# RORγ Inverse Agonism Redirects Tumour-Conditioned Emergency Myelopoiesis and Attenuates Established Tumour Progression

**DOI:** 10.64898/2026.10.01.755741

**Authors:** Iqraa Dhoparee-Doomah, Kim Ma, Ariane Brassard, Sabrina Leo, Shayan Hajhashemi, Yichen Li, Katy Milne, Ming Zi Xu, Qian Qiu, Manal Al Dow, Ahmed Fouda, France Bourdeau, Benjamin Gordon, Betty Giannias, Brad H Nelson, Jonathan Spicer, Lorenzo Ferri, Jean Tchervenkov, Jonathan Cools-Lartigue

**Author notes:** **Corresponding author**: Jonathan Cools-Lartigue, MD PhD Professor of Surgery, Division of Thoracic Surgery. These authors contributed equally and share first co-authorship (Iqraa Dhoparee-Doomah, Kim Ma).

## Abstract

Systemic inflammation is associated with poor outcomes in gastroesophageal adenocarcinoma (GEA), but how it contributes to disease progression remains unclear. We investigated retinoic acid receptor-related orphan receptor gamma (RORγ) as a regulator of tumour-conditioned myeloid responses. In treatment-naïve GEA patients, elevated circulating neutrophil-to-lymphocyte ratio (cNLR) tracked with circulating RORγ⁺ neutrophils and tumoural spatial organization of RORγ⁺ myeloid cells, rather than neutrophil abundance. In syngeneic tumour models, pharmacologic RORγ inverse agonism altered marrow progenitor composition before tumour-growth divergence and remodeled myeloid organization across bone marrow, tumour and lung despite no endpoint change in circulating neutrophil proportions, which did not capture the earlier marrow response or subsequent tumour trajectory. RORγ perturbation enhanced longitudinal tumour control with PD-L1 blockade, whereas effects on metastatic establishment depended on tumour conditioning and treatment timing. These findings identify RORγ-associated myeloid programming as a therapeutically tractable, context-dependent regulator linking systemic inflammation, hematopoiesis, tissue immune organization, and cancer progression.

**STATEMENT OF SIGNIFICANCE:** These findings establish RORγ-associated myeloid programming as a therapeutically tractable regulator linking systemic inflammation with tumour-conditioned myelopoiesis and immune organization in gastroesophageal adenocarcinoma. Targeting this axis enhanced PD-L1 blockade and modulated emergency myelopoiesis, while metastatic effects depended on tumour conditioning and treatment timing, highlighting critical principles for biomarker-guided therapeutic intervention.

## BACKGROUND

Cancer progression is shaped by systemic host immune responses and tumour-intrinsic biology (1). Tumour-derived inflammatory signals can alter hematopoiesis, expand immunoregulatory myeloid populations and remodel tissue immune-cell organization (2–4). Understanding these responses may identify therapeutic opportunities beyond tumour-directed approaches.

An elevated circulating neutrophil-to-lymphocyte ratio (cNLR) is associated with poorer outcomes across solid malignancies, including gastroesophageal adenocarcinoma (GEA) (5–9). However, cNLR does not capture neutrophil or lymphocyte functional diversity, developmental regulation, or tissue organization. Its prognostic association therefore raises the question of which immune processes accompany elevated cNLR and whether they can be therapeutically modified.

Neutrophils exhibit substantial functional plasticity and can support both antitumour and protumour immunity (10–12). Consequently, enrichment of a neutrophil phenotype in patients with elevated cNLR doesn’t establish a uniformly tumour-promoting role, as function may depend on developmental state and tissue microenvironment (3, 12, 13). Neutrophil plasticity and their role in host defence complicate broad depletion of mature neutrophils (10, 14, 15). Regulatory programmes governing tumour-associated emergency myelopoiesis (EM) therefore offer an upstream point of intervention that could change myeloid output without requiring broad depletion of mature neutrophils.

Retinoic acid receptor-related orphan receptor gamma (RORγ), encoded by *RORC*, is a candidate regulator of these programmes. Strauss and colleagues demonstrated that hematopoietic disruption of RORC1 impaired tumour-associated EM and reduced tumour burden (3). More recently, cholesterol-dependent RORγ activity was linked to expansion of suppressive myeloid populations, while pharmacological RORγ inhibition enhanced the efficacy of anti–PD-1 treatment in preclinical models (13).

*RORC* encodes RORγ (*RORC1*) and RORγt (*RORC2*), which differ at their amino termini but share DNA-binding and ligand-binding domains (16, 17). RORγt regulates type-17 differentiation and IL-17 production (16–18), pathways implicated in granulopoiesis, and metastatic progression (19–22). RORγ-associated biology therefore encompasses both myeloid developmental regulation and inflammatory signalling. We use “RORγ-associated” to describe the protein-level and pharmacological findings reported here, reserving “RORC1” for isoform-specific transcriptional or genetic evidence. How circulating RORγ-associated phenotypes relate to systemic inflammation, tissue organization and clinical outcome remains incompletely defined. It is unclear whether RORγ expression identifies a consistently adverse neutrophil phenotype or reflects a broader myeloid programme with consequences that vary across biological settings. We therefore asked whether perturbing RORγ-associated signalling alters tumour progression and the accompanying myeloid phenotype.

Here, we combined circulating and spatial analyses in GEA patients with pharmacological perturbation in mouse tumour models. We first examined relationships among RORγ-associated phenotypes, cNLR and survival, then assessed how RORγ inverse agonism affected tumour progression, marrow progenitors and tissue immune organization. We further evaluated therapeutic effects under different tumour-conditioning and intervention schedules and in combination with PD-L1 blockade. These analyses identified context-dependent clinical associations of RORγ-associated tissue features and contrasting effects of pathway inhibition on pulmonary metastasis, highlighting the importance of biological context and timing when targeting this pathway.

## RESULTS

### Circulating RORγ⁺ neutrophils are enriched in patients with high cNLR

Given the previously described role of RORC1 in tumour-associated EM (3), we first asked whether elevated cNLR was accompanied by enrichment of RORγ-expressing neutrophils. Among 28 treatment-naïve GEA patients, the proportion of circulating CD66b⁺CD16⁺ neutrophils expressing RORγ was higher in cNLR-high (≥4; n=8; median 14.7%, IQR 11.5–20.6%) compared to cNLR-low (<4; n=20; median 3.26%, IQR 1.15–7.86%; P=0.0039; **Fig. 1A–B**) patients.

**Figure 1:**
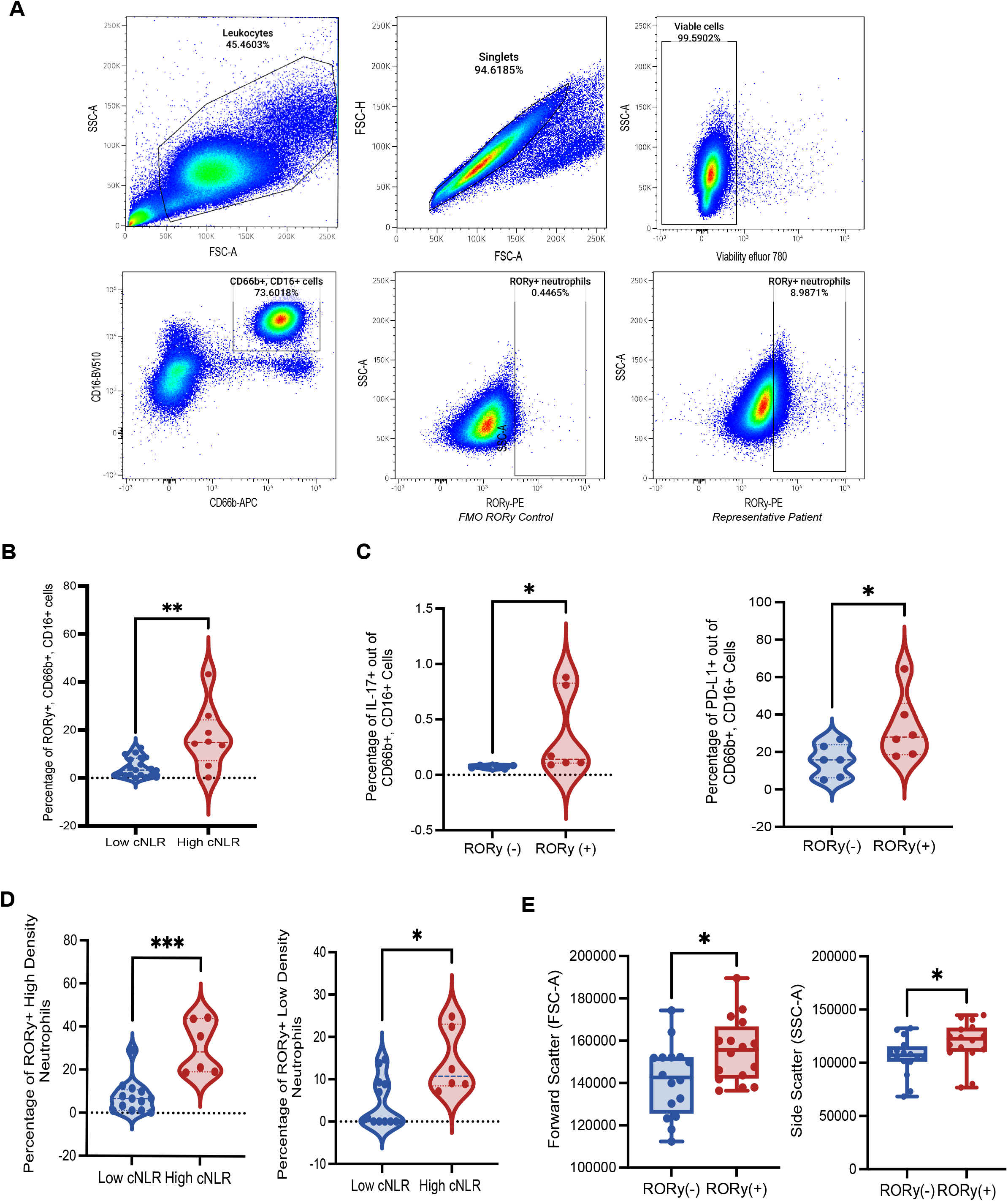
Circulating RORγ⁺ neutrophils are enriched in patients with high cNLR. (A) Representative gating strategy used to identify circulating CD66b⁺CD16⁺RORγ⁺ neutrophils, including the RORγ FMO control. (B) Frequency of circulating RORγ⁺ neutrophils in patients with low versus high cNLR. (C) IL-17 and PD-L1 expression in RORγ⁻ and RORγ⁺ neutrophils. (D) Frequency of RORγ⁺ cells within high-density and low-density neutrophil fractions. (E) Forward- and side-scatter characteristics reflecting cell size and granularity. *P<0.05, **P<0.01, ***P<0.001.

We next characterized the circulating RORγ⁺ phenotype. In paired patient analyses, IL-17⁺ and PD-L1⁺ neutrophils were higher for the RORγ⁺ than the RORγ⁻ phenotype (n=6; P=0.0312 for each comparison; **Fig. 1C**). RORγ enrichment in patients with high cNLR was evident in both high-density neutrophils (n=6 high versus 13 low; P=0.0005) and low-density neutrophils (n=6 high versus 13 low; P=0.0139; **Fig. 1D**). Within individual patients, RORγ⁺ neutrophils also exhibited higher forward and side scatter than RORγ⁻ neutrophils (n=16; P<0.05 for both; **Fig. 1E**), consistent with differences in relative cell size and internal complexity.

These findings linked elevated cNLR to enrichment of a phenotypically distinct RORγ⁺ neutrophil population across both density fractions but did not establish whether RORγ marked a tumour-promoting population or a broader myeloid programme. We therefore examined the anatomical distribution and clinical associations of RORγ-associated phenotypes in human tissues.

### Spatial organization of RORγ-associated tissue phenotypes shows distinct relationships with systemic inflammation and survival

We examined a GEA tissue microarray (TMA) comprising 29 patients, with samples from histologically normal esophagus and stomach, tumour periphery and tumour core. Clinicopathological features are summarized in **Supplementary Table S1**. Multiplex immunofluorescence was used to identify the ARG1⁺ tissue neutrophil compartment and assess RORγ and IL-17 expression and localization (**Fig. 2A**). Abundance varied across anatomical compartments: the median patient-level ARG1⁺ cell count per core was 4.0 in normal oesophagus, 25.5 in normal stomach, 33.5 at the tumour periphery and 58.0 in the tumour core. ARG1⁺ cells were more abundant in normal stomach than in paired normal oesophagus samples (n=28; q=0.00090) and in tumour core than in paired normal stomach samples (n=29; q=0.018; **Fig. 2B**).

**Figure 2:**
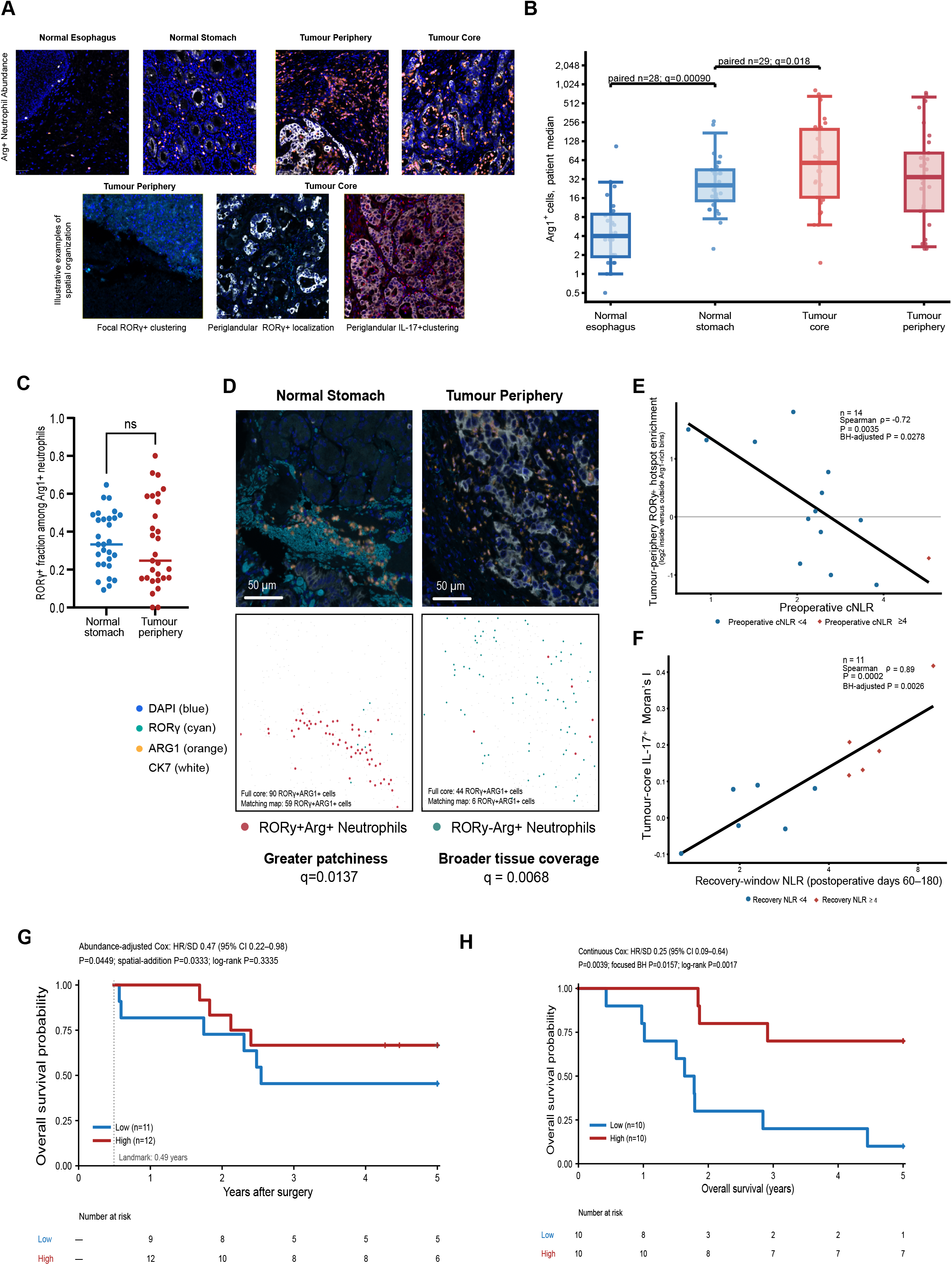
Spatial organization of RORγ-associated tissue phenotypes shows distinct relationships with systemic inflammation and survival. (A) Representative microscopy of normal esophagus, normal stomach, tumour core, and tumour periphery. (B) Arg1+ cell abundance across gastroesophageal compartments. (C) Prevalence of RORγ⁺ Arg1+ neutrophils in normal stomach and tumour periphery. (D) Spatial distribution of RORγ⁺ neutrophils in normal stomach and tumour periphery. (E) Association between tumour-periphery RORγ⁺ Arg1+ hotspot enrichment and preoperative cNLR. (F) Association between tumour-core IL-17+ Moran’s I and recovery-window cNLR. (G) Association between tumour-periphery RORγ⁺ Arg1+ hotspot enrichment and overall survival. (H) Association between tumour-core RORγ⁺IL-17⁺ARG1⁺ cells proximity to CK7+ epithelial cells and overall survival. Proximity threshold was defined as within 20 µm. Continuous Cox models are primary; Kaplan–Meier curves are illustrative. **Abbreviations**: RORγ, retinoic acid receptor-related orphan receptor-γ; Arg-1: Arginase 1; cNLR: circulating neutrophil-to-lymphocyte ratio; CK7: cytokeratin 7

We next distinguished population composition from spatial distribution. The fraction of ARG1⁺ cells expressing RORγ did not differ significantly between normal stomach and tumour periphery (median, 33.3% versus 24.8%; paired n=29; P=0.611; **Fig. 2C**). Their spatial distribution nevertheless differed. Among 14 paired patients meeting spatial-analysis criteria, RORγ⁺ARG1⁺ cells occupied a greater fraction of eligible tissue tiles at the tumour periphery (q=0.0068), whereas normal stomach showed greater variability in their prevalence between tiles (q=0.0137; **Fig. 2D**). Thus, the tumour periphery showed broader tissue coverage, while normal stomach showed a patchier distribution, despite no detectable difference in RORγ⁺ composition.

We then asked whether specific spatial arrangements were associated with systemic inflammation. Greater tumour-periphery RORγ⁺ARG1⁺ hotspot enrichment, measuring preferential localization within ARG1-rich regions, was associated with lower preoperative cNLR (n=14; Spearman ρ=−0.72; P=0.0035; q=0.0278; **Fig. 2E**). A different relationship was observed in the tumour core: greater spatial clustering of IL-17⁺ cells, measured by Moran’s I, was associated with higher recovery cNLR measured during postoperative years 0.16–0.49 (n=11; Spearman ρ=0.89; P=0.00023; q=0.0026; **Fig. 2F**). We next examined whether these spatial features carried prognostic information.

Greater tumour-periphery RORγ⁺ARG1⁺ hotspot enrichment was associated with better overall survival (OS) after adjustment for matched cell abundance. Among 23 patients surviving to the postoperative landmark at 0.49 years, with follow-up capped at five years after surgery, the hazard ratio per standard-deviation increase in enrichment was 0.47 (95% CI, 0.22–0.98; P=0.045; 10 deaths; **Fig. 2G**). Adding hotspot enrichment improved model fit relative to abundance alone (likelihood-ratio P=0.033). Apparent discrimination also increased, with Harrell’s C-index rising from 0.57 to 0.73 and the five-year time-dependent AUC from 0.73 to 0.83. These descriptive estimates were obtained in the same cohort used to fit the models. Greater peripheral enrichment was therefore associated with both lower preoperative cNLR and better survival. The landmark model included ten deaths and adjusted for cell abundance. Stage and treatment were not included as covariates.

Tumour-core IL-17⁺ clustering did not show a clear association with survival through five years after surgery (n=24; 12 deaths; HR per standard deviation, 1.19; 95% CI, 0.65–2.21; P=0.571). An exploratory post hoc analysis separating follow-up into 0–2 and 2–5 years suggested variation in the association over time (nominal interaction P=0.023; **Supplementary Fig. S1**). The estimated association was adverse during the first two years and favourable among patients surviving beyond two years, although both interval-specific confidence intervals included the null and each interval contained only six deaths.

A further survival association involved proximity to the tumour epithelium. A greater proportion of tumour-core RORγ⁺IL-17⁺ARG1⁺ cells located within 20 µm of CK7⁺ epithelial cells was associated with better OS, with follow-up capped at five years (n=20; 12 deaths; HR per standard deviation, 0.25; 95% CI, 0.09–0.64; P=0.0039; q=0.0157 within the focused four-feature analysis; **Fig. 2H**). No clear association between this proximity feature and NLR was identified.

Together, these observations distinguished tissue features associated with systemic inflammation from those associated with survival. Peripheral hotspot enrichment was related to both, whereas tumour-core IL-17⁺ clustering and epithelial proximity showed different clinical associations. These findings motivated testing whether perturbation of RORγ-associated signalling could alter tumour progression and its accompanying host phenotype.

### RORγ inverse agonism reduces intratumoural RORγ⁺ neutrophil frequency and pulmonary metastatic burden

We investigated whether pharmacological RORγ inhibition affected tumour progression and neutrophil populations in syngeneic tumour models. RORC expression was minimal or undetectable in YTN16 gastric, MC38 colorectal and H59 lung adenocarcinoma cells, and TF-S10 didn’t substantially impair tumour-cell viability, clonogenic growth or migration across the concentrations tested (**Supplementary Fig. S2-3**). These findings supported investigating a contribution from the host compartment to the effects of RORγ inhibition.

We evaluated TF-S10 in established syngeneic tumours, initiating treatment seven days after implantation (**Fig. 3A**). YTN16 tumour volumes were numerically lower in treated mice, although the longitudinal treatment × time interaction did not reach statistical significance (Greenhouse–Geisser-corrected P = 0.0678; **Fig. 3B**). Independent experiments in MC38-RFP and H59-GFP models also showed lower tumour growth with TF-S10 treatment (**Supplementary Fig. S4**).

**Figure 3:**
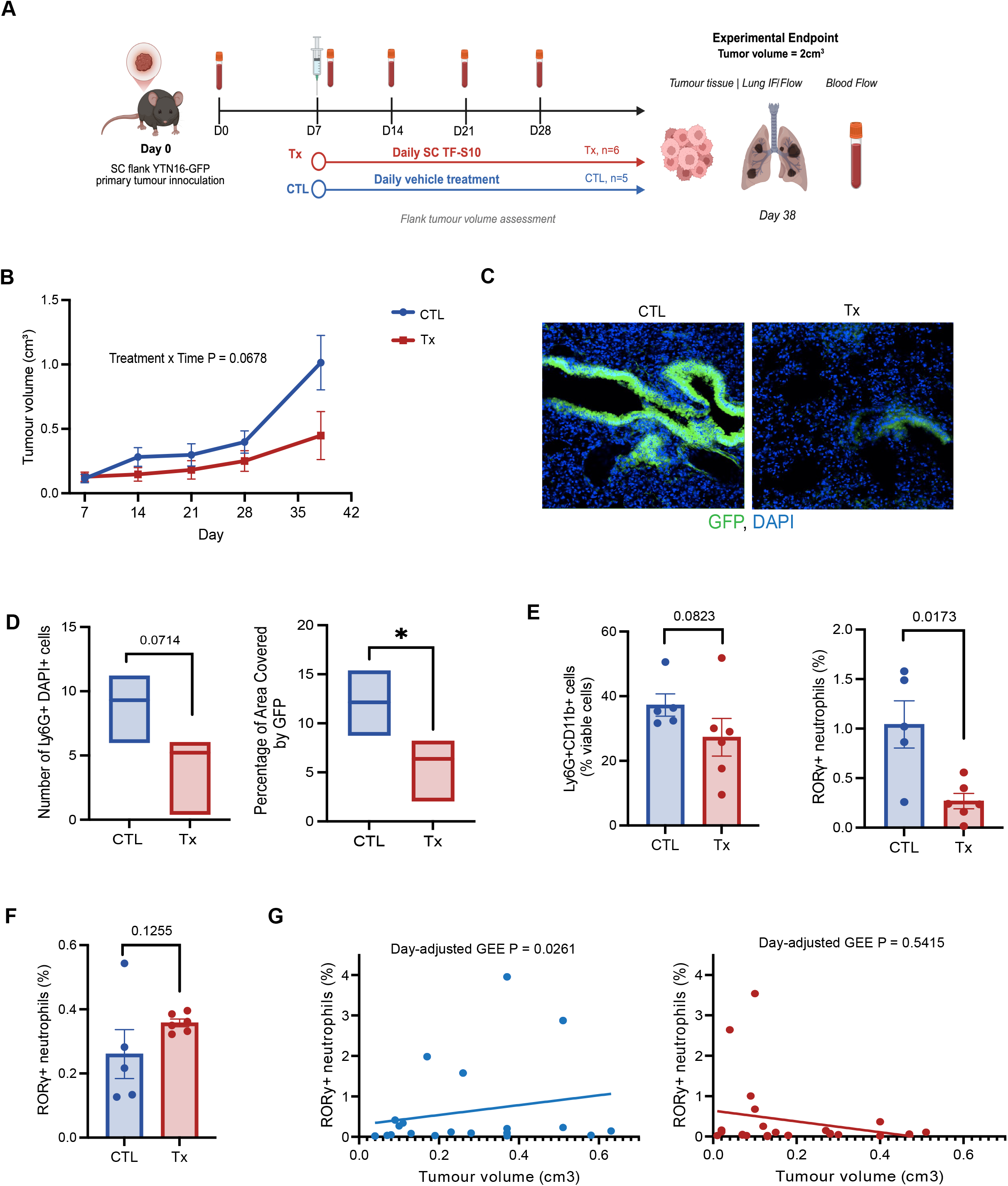
RORγ inverse agonism reduces intratumoural RORγ+ neutrophil frequency and pulmonary metastatic burden. (A) Experimental design of the established YTN16-GFP tumour model with 25mg/kg TF-S10 treatment initiated on D7. (B) Longitudinal tumour growth in control (CTL) and TF-S10-treated (Tx) mice. (C) Representative pulmonary GFP⁺ metastases. (D) GFP⁺ metastatic burden and pulmonary Ly6G⁺ neutrophil infiltration. (E) Intratumoural Ly6G⁺ neutrophils and RORγ⁺ neutrophils at endpoint. (F) Circulating RORγ⁺ neutrophils at endpoint. (G) Association between tumour volume and RORγ⁺ neutrophil frequency according to treatment. CTL, control; Tx, TF-S10 treatment; RORγ, retinoic acid receptor-related orphan receptor-γ; GFP, green fluorescent protein; Ly6G, lymphocyte antigen 6 complex locus G6D. *P<0.05; **P<0.01. Exact P values are shown where indicated.

In the YTN16-GFP model, pulmonary metastatic burden was lower following TF-S10 treatment. In lung specimens from three control and five treated mice, GFP⁺ tumour burden, quantified as the percentage of image area covered by GFP and summarized within each mouse, was lower in the treated group (P=0.0357; **Fig. 3C-D**). Pulmonary Ly6G⁺ neutrophil counts tended to be lower, although the difference was not statistically significant (P=0.0714; **Fig. 3D**). The reduction in pulmonary burden was detected despite no statistically significant separation in primary-tumour growth trajectories.

We next examined intratumoural and circulating neutrophil populations. Within YTN16 tumours, the proportion of Ly6G⁺CD11b⁺ neutrophils among viable cells was numerically lower following treatment, but the difference was not statistically significant (P = 0.0823; **Fig. 3E**). By contrast, the proportion of intratumoural neutrophils expressing RORγ was significantly lower in treated mice (P = 0.0173; **Fig. 3E**). At the experimental endpoint, the proportion of circulating neutrophils expressing RORγ did not differ significantly between groups (P = 0.1255; **Fig. 3F**).

We additionally explored the relationship between tumour volume and circulating RORγ⁺ neutrophil frequency over time. In a generalized estimating equation model accounting for repeated measurements within mice and adjusting for sampling day, tumour volume was positively associated with circulating RORγ⁺ neutrophil frequency in control mice (P = 0.0261; **Fig. 3G**). No statistically significant association was detected in treated mice (P = 0.5415; **Fig. 3G**). However, the tumour volume × treatment interaction was not statistically significant (P = 0.0613), so these exploratory findings did not establish that treatment altered the association.

### RORγ inverse agonism alters marrow progenitor composition before detectable tumour-growth inhibition

Because tumour suppression was not accompanied by a detectable reduction in circulating RORγ⁺ neutrophil frequency, we next asked whether TF-S10 altered the upstream marrow progenitor compartment and when these changes emerged relative to tumour control. We used the subcutaneous MC38-RFP model, which generates primary flank tumours without overt metastasis over the experimental interval. Treatment began seven days after tumour implantation, with independent terminal cohorts sampled during tumour progression and a separate cohort assessed at a fixed D28 endpoint (**Fig. 4A**).

**Figure 4:**
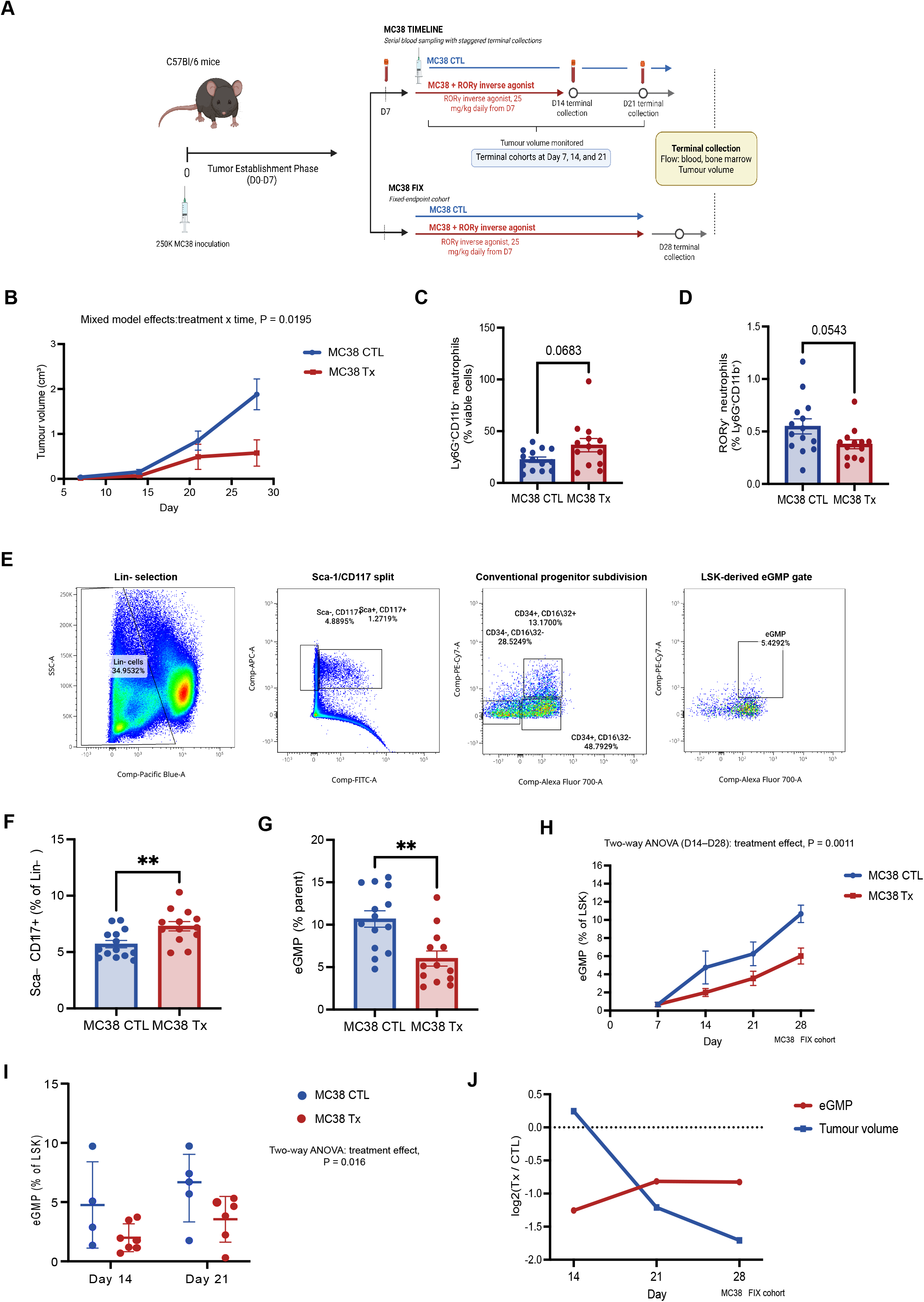
RORγ inverse agonism alters marrow progenitor composition before detectable tumour-growth inhibition. (A) Experimental design. Syngeneic MC38-RFP colorectal tumours were established subcutaneously in C57BL/6 mice in two separate experiments: one with a fixed endpoint D28 (MC38 FIX) and the other with endpoints at D7, D14 and D21 (MC38 Timeline). RORγ inverse agonist TF-S10 (Tx) and vehicle (CTL) treatment commenced on D7 after tumour implantation and continued until experimental endpoint. (B) Tumour progression under different treatment conditions in the MC38 FIX cohort. (C-D) Prevalence of circulating (C) neutrophils and (D) RORγ+ neutrophils at experimental endpoint. (E) Flow cytometric gating strategy for BM progenitor populations. (F-G) Proportion of (F) Sca-, CD117+ conventional myeloid progenitors and (G) emergency granulocyte–monocyte progenitor (eGMP) under different treatment conditions in the MC38 FIX cohort at endpoint. (H) Prevalence of eGMP throughout disease progression under different treatment conditions. (I) Proportion of eGMPs between D14 and D21 post-tumour inoculation under different treatment conditions. (J) Relationship between tumour volume and prevalence of eGMPs. **P<0.01. Exact P values are shown where indicated. **Abbreviations**: CTL: vehicle control; Tx: TF-S10 treatment; RORγ, retinoic acid receptor-related orphan receptor-γ; eGMP: emergency granulocyte–monocyte progenitor

TF-S10 suppressed primary tumour growth at later time points (P=0.0195; **Fig. 4B**). At the D28 endpoint, however, neither the frequency of circulating Ly6G⁺CD11b⁺ neutrophils among viable cells nor the proportion of these neutrophils expressing RORγ differed significantly between control and treated mice (P=0.0683 and P=0.0543, respectively; **Fig. 4C–D**). Thus, tumour control was again observed without a statistically significant reduction in the circulating RORγ⁺ neutrophil fraction.

We then examined the bone marrow (BM) progenitor compartment. Flow cytometry distinguished conventional Sca-1⁻CD117⁺ myeloid progenitors within the lineage-negative population from emergency granulocyte–monocyte progenitors (eGMPs) within the lineage-negative Sca-1⁺CD117⁺ compartment (LSK; **Fig. 4E**). At D28, TF-S10 increased the fraction of Sca-1⁻CD117⁺ progenitors among lineage-negative cells and reduced eGMP frequency within the LSK compartment (both P<0.01; **Fig. 4F–G**). These findings demonstrated different treatment-associated changes in the representation of the two progenitor populations.

The reduction in eGMP frequency was evident early after treatment initiation. On D14, following seven days of TF-S10, eGMPs constituted 1.99% of LSK cells in treated mice compared with 4.76% in controls, a numerical reduction of more than half. Their frequency remained lower in treated mice on D21 (3.55% versus 6.26%). Analysis of these independent D14 and D21 cohorts identified an overall treatment effect on eGMP frequency (two-way ANOVA, P=0.016; **Fig. 4I**). By D28, eGMP frequency remained lower in treated mice (6.02% versus 10.67%; Mann–Whitney P=0.0027; **Fig. 4H**). These individual time-point estimates are descriptive; the ANOVA tests an overall early-cohort treatment effect. Comparison with tumour-volume trajectories showed that the D14 eGMP difference preceded the later separation of tumour growth between treatment groups **(Fig. 4J**).

These findings identified an early marrow progenitor response to RORγ inverse agonism that preceded detectable tumour-growth inhibition. This temporal ordering supports an effect on tumour-associated haematopoietic programming, although it does not establish that the progenitor changes mediated subsequent tumour control. We next asked whether the consequences of RORγ perturbation depended on prior tumour conditioning and the timing of intervention.

### Tumour conditioning and intervention timing shape the effect of RORγ inverse agonism on pulmonary metastatic burden

The early marrow changes observed following RORγ inverse agonism raised the question of whether treatment efficacy depended on the host context in which inhibition occurred. We therefore examined whether prior tumour conditioning and the timing of TF-S10 administration influenced pulmonary metastatic burden following intravenous tumour-cell challenge. MC38-RFP cells were administered either to mice without a pre-existing tumour or to mice bearing a subcutaneous MC38-RFP flank tumour, which provided tumour conditioning without overt spontaneous metastasis over the experimental interval (**Fig. 5A**).

**Figure 5:**
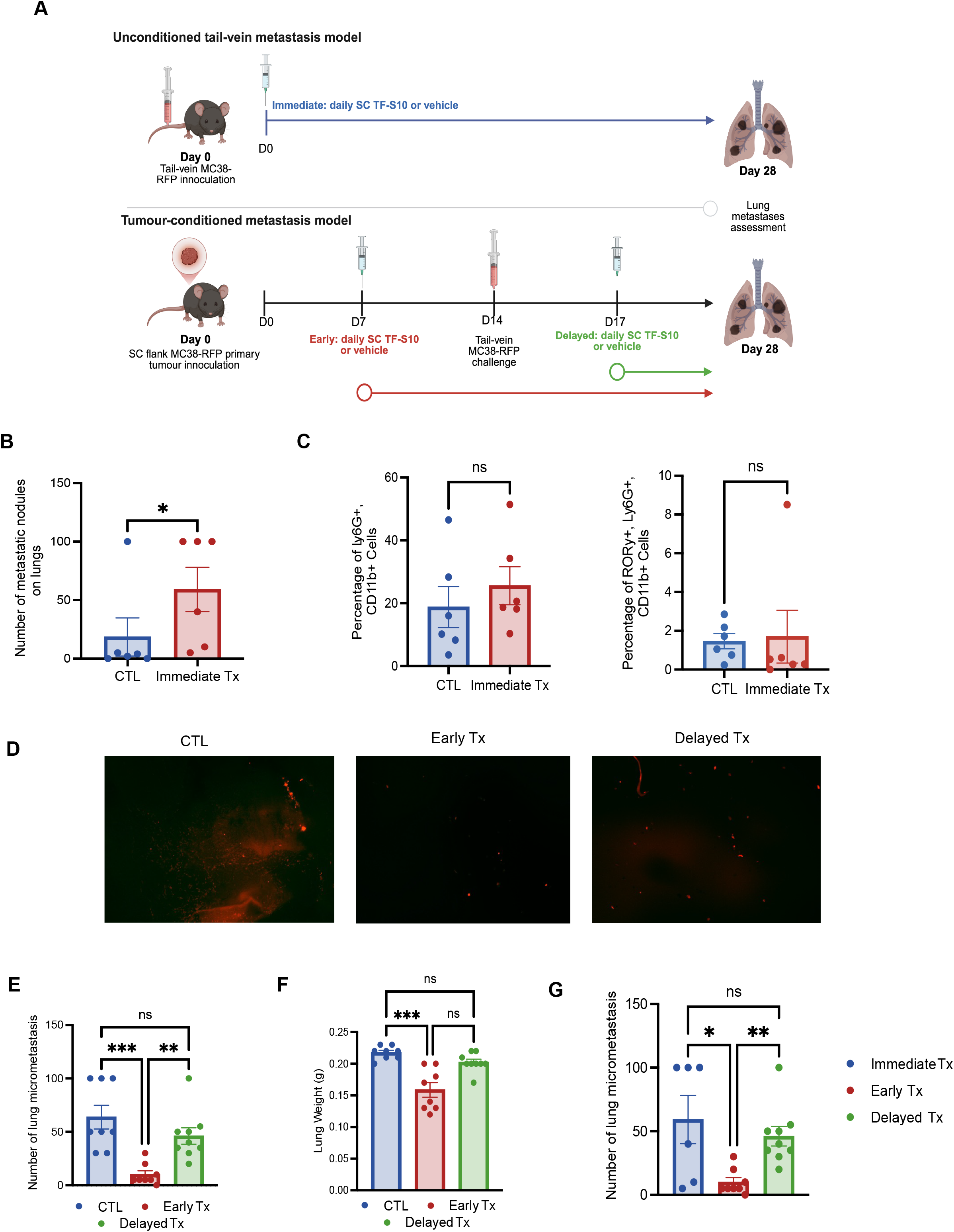
Tumour conditioning and intervention timing shape the effect of RORγ inverse agonism on pulmonary metastatic burden. (A) Experimental design of intravenous (IV) tumour cell inoculum without conditioning primary tumour and with conditioning primary tumour. (B) Number of lung metastatic nodules under different treatment conditions in unconditioned tail-vein metastasis model. (C) Circulating neutrophils and RORγ+ neutrophils prevalence in unconditioned tail-vein metastasis model. (D) Representative field (4x magnification) of mouse lung showing micrometastasis in red (RFP-positive) in tumour-conditioned metastasis model. (E) Number of lung micrometastasis and (F) lung weight in tumour-conditioned metastasis model under different treatment conditions. (G) Exploratory analysis of treatment timing across all three metastasis cohorts. **Abbreviations:** CTL: vehicle control; Tx: TF-S10 treatment; RORγ, retinoic acid receptor-related orphan receptor-γ; SC = subcutaneous

In mice without prior tumour conditioning, TF-S10 began 24 hours after intravenous MC38-RFP challenge; lung nodules were assessed on D28. Under these conditions, TF-S10 increased pulmonary nodule burden: vehicle-treated mice developed a median of 3 nodules, compared with 70 in treated mice (n=6/group; exact P=0.039; **Fig. 5B**). The response was heterogeneous: three of six treated mice reached the reporting ceiling of 100 nodules, whereas two treated mice had fewer than 10 nodules; one control mouse also reached the 100-nodule ceiling. No technical or dosing deviation was identified that accounted for these discordant animals. Despite inter-animal variability and the upper reporting limit, these data demonstrate that TF-S10 increased pulmonary metastatic burden in this setting.

This increase in pulmonary metastatic burden was not accompanied by a statistically significant difference in the frequency of circulating Ly6G⁺CD11b⁺ neutrophils or the proportion of these neutrophils expressing RORγ at endpoint (exact P=0.240 for each comparison; **Fig. 5C**).

We next examined treatment timing in tumour-conditioned mice. Following subcutaneous flank-tumour implantation on D0, mice were challenged intravenously with MC38-RFP cells on D14. TF-S10 was initiated either on D7, before intravenous challenge, or on D17, after challenge; pulmonary burden was assessed on D28 (**Fig. 5A**). Early treatment markedly reduced the median pulmonary nodule count from 50 in controls to 5 with TF-S10 (n=8/group; Dunn–Bonferroni-adjusted P=0.0008; **Fig. 5D-E**). Mice receiving delayed treatment developed a median of 40 nodules (n=9), significantly more than the early-treatment group, with no significant difference from controls (Dunn–Bonferroni-adjusted P=0.0073 and P=1.000, respectively; **Fig. 5D-E**).

Lung weight provided a complementary burden-related endpoint. Early treatment reduced median lung weight from 0.220 g in controls to 0.155 g (Dunn–Bonferroni-adjusted P=0.0003; **Fig. 5F**). Median lung weight following delayed treatment was 0.200 g, with no significant difference from controls after adjustment for multiple comparisons (P=0.2391; **Fig. 5F**). These endpoint measurements did not establish a change in metastatic growth rate.

Across these experimental conditions, the direction and magnitude of the response to TF-S10 differed with tumour-conditioning history and treatment schedule (**Fig. 5G**). Treatment increased pulmonary metastatic burden in mice without a pre-existing tumour, whereas the clearest protective effect occurred in tumour-bearing mice treated before intravenous challenge. Delayed treatment produced a weaker, endpoint-dependent signal. Because the protocols also differed in treatment duration and the interval between intravenous challenge and assessment, these findings did not define a discrete mechanistic window or distinguish effects on initial metastatic establishment from subsequent outgrowth. They nevertheless supported examining RORγ inhibition within the broader tumour–host context and motivated further investigation of myeloid tissue organization

### Exploratory spatial profiling identifies compartment-specific myeloid changes following RORγ inverse agonism

The human analyses identified distinct relationships between myeloid tissue organization, systemic inflammation and survival, highlighting the importance of examining cellular phenotypes within their anatomical context. We therefore extended this spatial approach to the murine experiments, asking whether the tumour and marrow responses to RORγ inverse agonism were accompanied by tissue changes not apparent from circulating RORγ⁺ neutrophil frequency alone. Exploratory spatial profiling was performed on primary tumours, histologically uninvolved lung and BM collected at D28 from TF-S10-treated and control MC38-RFP-bearing mice. Independent cohorts provided three animals per treatment group for tumour analysis, three per group for lung analysis and two specimens per group for marrow analysis (**Fig. 6A**). Multiple predefined fields or tiles were sampled within each tissue. The mouse or specimen remained the biological replicate, with nested analyses used to characterize variation across the sampled regions.

**Figure 6:**
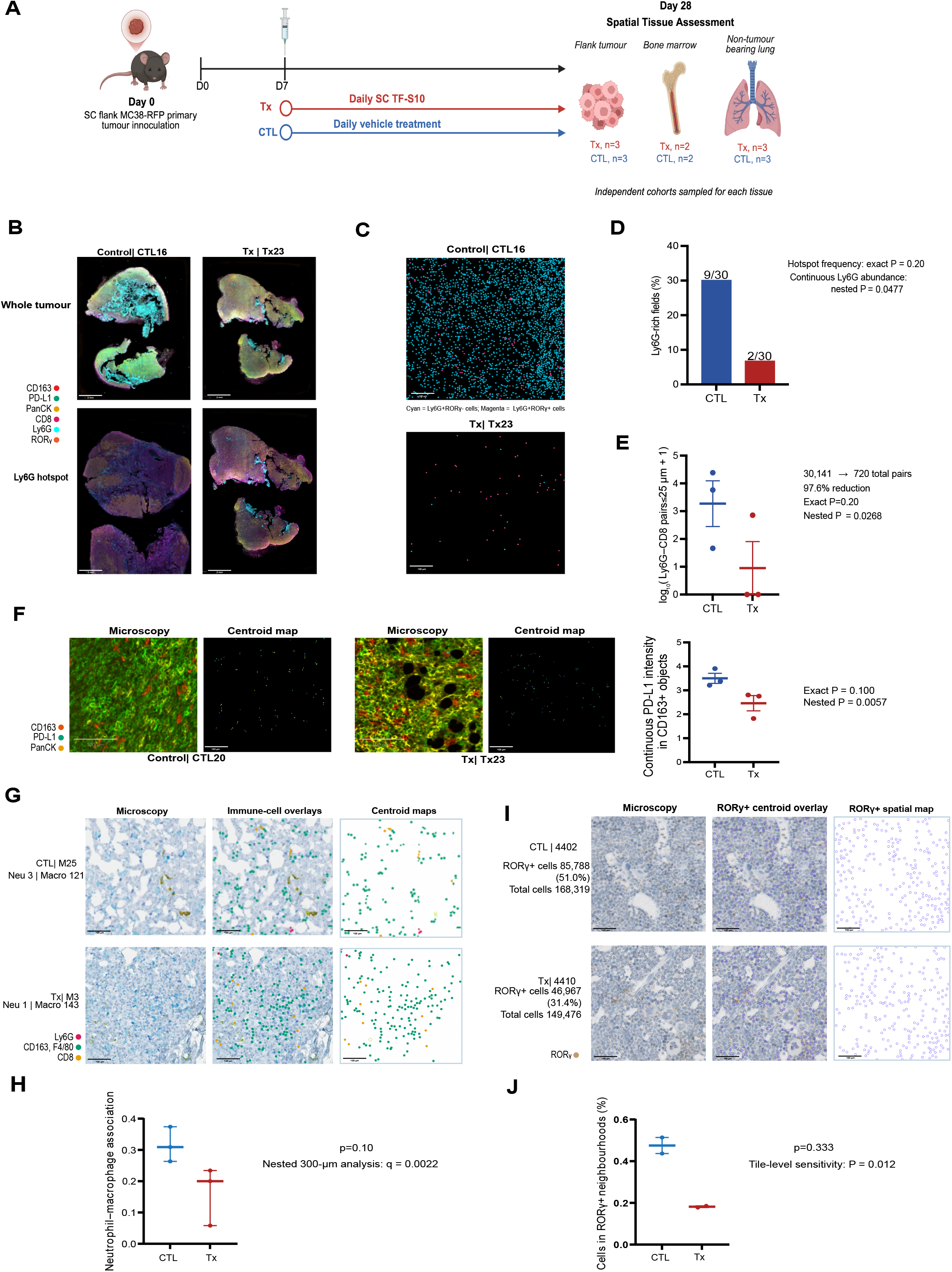
Exploratory spatial profiling identifies compartment-specific myeloid changes following RORγ inverse agonism. (A) Experimental design for cross compartment spatial analysis. (B) Representative whole-tumour and Ly6G hotspot multiplex immunofluorescence images. Fluorescence channels: PanCK (yellow), CD163 (red), PD-L1 (green), Ly6G (cyan), CD8 (magenta) and RORγ (orange). (C) Representative centroid maps of Ly6G⁺ neutrophils classified according to RORγ expression. Cyan denotes Ly6G⁺RORγ⁻ cells and magenta denotes Ly6G⁺RORγ⁺ cells. (D) Quantification of the frequency of Ly6G-rich hotspot fields, defined as sampled tumour fields in which Ly6G⁺ cells comprised >1% of all evaluable cells carrying a binary Ly6G call. (E) Number of Ly6G+-CD8+ spatial pairs within 25 µm in CTL and Tx tumours. (F) Multiplex immunofluorescence and centroid maps of representative tumour field and quantification of continuous PD-L1 intensity in CD163⁺ myeloid cells. Fluorescence channels: PanCK yellow; CD163 red; PD-L1 green. In the centroid maps, each dot represents the centroid of a CD163⁺ cell and is coloured according to the mean whole-cell PD-L1 intensity using a shared fixed colour scale (0.7846–5.9566 a.u.), with colours ranging from purple through blue, teal, and green to yellow, corresponding to increasing PD-L1 expression. (G) Representative maps of regional neutrophil–macrophage organization in histologically uninvolved lung. Macrophage (F4/80, CD163): blue (microscopy) and green (centroid); neutrophils (Ly6G): purple (microscopy) and magenta (centroid); CD8: Dab (microscopy) and yellow (centroid). (H) Quantification of regional neutrophil–macrophage association in lung at the mouse level and across prespecified 300-µm fields with tiles nested within each mouse. (I) Representative BM maps illustrating local RORγ⁺ cell organization and quantification of the proportion of cells occupying RORγ⁺-enriched local neighbourhoods (≥50% RORγ⁺ cells). (J) Tile-based sensitivity analysis of local RORγ⁺ cell representation within BM. Primary treatment comparisons used mouse or specimen as the biological replicate, with mixed-effects analyses of nested tumour fields or tissue tiles performed as supporting analyses of within-tissue spatial organization. **Abbreviations**: CTL: vehicle control; Tx: TF-S10 treated; RORγ, retinoic acid receptor-related orphan receptor-γ; SC = subcutaneous

Within primary tumours, microscopy and cell-position maps illustrated differences in Ly6G⁺ neutrophil distribution between control and treated specimens (**Fig. 6B–C**). Ly6G-rich fields were less frequent following TF-S10 treatment, occurring in 2 of 30 sampled fields compared with 9 of 30 control fields (mouse-level exact P=0.20; nested P=0.0476; **Fig. 6D**). Treated tumours also contained fewer Ly6G⁺–CD8⁺ spatial pairs within 25 µm (mouse-level exact P=0.20; nested P=0.0268; **Fig. 6E**). These findings described reduced regional neutrophil enrichment and fewer observed neutrophil–CD8⁺ cell pairs; the pair-count reduction could also reflect the lower representation of neutrophils.

We additionally examined the PD-L1 phenotype of tumour-associated CD163⁺ cells. Continuous PD-L1 intensity within these cells was lower in treated specimens (mouse-level exact P=0.10; nested P=0.0057; **Fig. 6F**). This observation suggested that the tissue response to TF-S10 included a change in a checkpoint-associated myeloid phenotype alongside changes in neutrophil distribution.

We next examined a distal tissue without overt tumour involvement. In histologically uninvolved lung, regional neutrophil–macrophage association was lower in treated animals (mean Spearman ρ, 0.164 versus 0.316 in controls; mouse-level P=0.10; **Fig. 6G–H**). An exploratory nested analysis of predefined 300-µm fields showed a concordant difference (β=−0.165; P=0.000446; BH q=0.00223). Thus, a treatment-associated difference in regional myeloid organization was also observed outside the primary tumour.

In BM, the proportion of cells located within neighbourhoods containing at least 50% RORγ⁺ cells decreased from 47.5% in control specimens to 18.2% in treated specimens (**Fig. 6I–J**). With two specimens per group, the specimen-level permutation test did not reach statistical significance (P=0.333). A complementary tile-based analysis showed lower local RORγ⁺ cell representation following treatment (29.7% versus 49.5%; tile-level P=0.012). This provided a supporting spatial readout rather than additional biological replication or a second test of the same neighbourhood endpoint.

These exploratory analyses identified compartment-specific differences in myeloid organization and phenotype following RORγ inverse agonism. The small independent cohorts limited inference at the biological-replicate level, while the regional analyses characterized the tissue features accompanying treatment. In particular, the change in PD-L1 intensity within tumour-associated CD163⁺ cells provided a rationale to test whether combining TF-S10 with PD-L1 blockade improved tumour control.

### RORγ inverse agonism enhances tumour control by PD-L1 blockade

We evaluated TF-S10, anti-PD-L1 and their combination in three independent cohorts of mice bearing subcutaneous MC38-RFP tumours, with five mice initially assigned to each treatment group. Cohorts 1 and 3 included corresponding control arms, whereas cohort 2 included active-treatment arms only (**Fig. 7A**). Treatment began seven days after tumour implantation, and tumour growth was followed through D28, when endpoint tumour burden and gross response categories were assessed.

**Figure 7:**
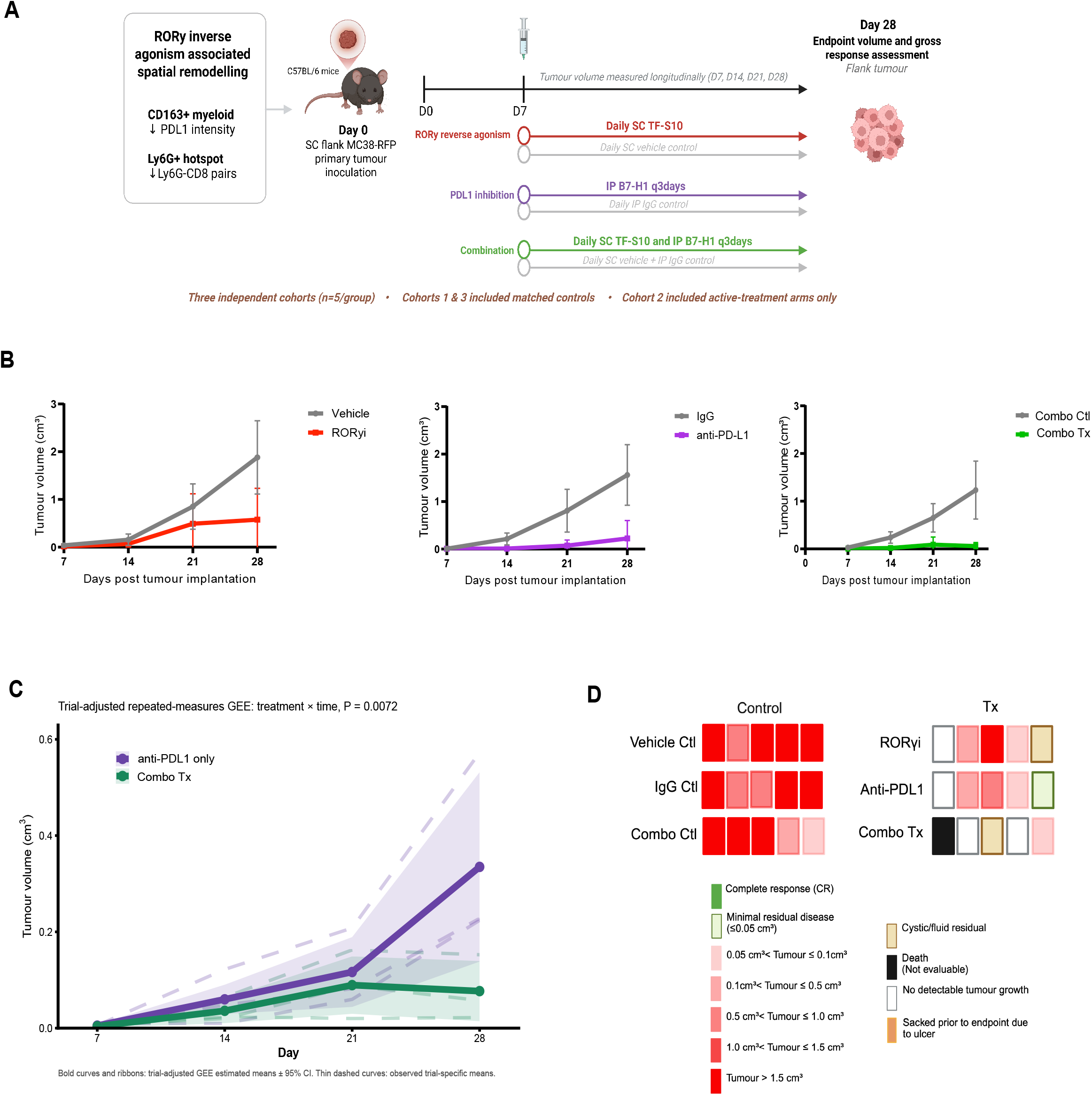
RORγ inverse agonism enhances tumour control by PD-L1 blockade. (A) Study design and treatment schedule across three independent cohorts of subcutaneous MC38-RFP tumour-bearing mice (n=5/group), treated with TF-S10 (RORγi), anti-PD-L1 (B7-H1) or combination of both (combo Tx). Cohorts 1 and 3 included corresponding control groups, whereas cohort 2 included active-treatment groups only. (B) Representative longitudinal tumour-growth trajectories in cohort 1 for controls, TF-S10, anti-PD-L1 and combo Tx groups. (C) Longitudinal tumour-growth trajectories across all three cohorts comparing anti-PD-L1 monotherapy with combo Tx. (D) Gross tumour response categories at experimental endpoint (D28) in cohort 1. **Abbreviations:** RORγi: TF-S10 only; Combo Ctl: Combination control (IgG + TF-S10 vehicle control); Combo Tx: combination treatment (anti-PDL1+ TF-S10); IgG: Immunoglobulin G; CR: complete response; SC = subcutaneous; IP: intraperitoneal

Representative trajectories from cohort 1 showed lower tumour growth with TF-S10, anti-PD-L1 and combination treatment than with their respective controls (**Fig. 7B**). Combination-treated mice showed sustained suppression of mean tumour volume through D28, with little additional growth between D21 and D28.

To assess whether adding TF-S10 improved tumour control beyond PD-L1 blockade alone, we compared longitudinal tumour-growth trajectories in the combined data from all three cohorts using a trial-adjusted repeated-measures generalized estimating equation model. Trajectories differed significantly between anti-PD-L1 monotherapy and combination treatment (treatment-by-time interaction, P=0.0072; **Fig. 7C**). Whereas estimated mean tumour volume increased during later follow-up with anti-PD-L1 alone, combination treatment maintained tumour-growth suppression through D28.

Gross response categories in cohort 1 provided a descriptive assessment of residual tumour burden (**Fig. 7D**). Control animals predominantly occupied the larger tumour-volume categories, while active-treatment groups included animals with no detectable tumour growth or low residual burden. The category “no detectable tumour growth” was assigned to one of five mice receiving TF-S10 alone and one of five receiving anti-PD-L1 alone. Among four evaluable combination-treated mice, two had no detectable tumour growth, one had a small solid residual tumour and one had a cystic/fluid residual lesion. The fifth combination-treated mouse died before the planned endpoint assessment and was not included in the evaluable endpoint-response denominator. These categories were descriptive and did not constitute a formal comparison of response rates.

Together, these findings demonstrated improved longitudinal tumour control when TF-S10 was added to PD-L1 blockade in the established MC38-RFP tumour setting. The combination benefit supports further investigation of RORγ-associated signalling as a therapeutic target alongside checkpoint blockade; whether the observed myeloid spatial and PD-L1 changes mediated this benefit remains unresolved.

## DISCUSSION

This study links RORγ-associated phenotypes to systemic inflammation and tissue organization in human GEA and demonstrates that pharmacological RORγ inverse agonism can alter tumour progression and the accompanying myeloid phenotype in mouse models. The human findings revealed distinct relationships among circulating RORγ⁺ neutrophils, anatomical context, cNLR and survival. In mice, treatment altered marrow progenitor composition before tumour-growth divergence, remodeled myeloid organization across tissues, and enhanced tumour control with PD-L1 blockade. However, effects on pulmonary metastatic burden varied with tumour conditioning and treatment timing. Together, these findings support RORγ-associated myeloid programming as a therapeutically modifiable component of the tumour–host response whose significance cannot be inferred from marker expression or endpoint circulating cell frequencies alone.

The human spatial analyses demonstrate the importance of distinguishing cellular abundance from tissue organization. Normal stomach and tumour periphery differed in the distribution of RORγ⁺ARG1⁺ cells despite no detectable difference in their proportion within the ARG1⁺ compartment. At the tumour periphery, greater RORγ⁺ARG1⁺ hotspot enrichment was associated with lower preoperative cNLR and better survival, including after adjustment for cell abundance, suggesting that spatial organization contained information beyond population size. Conversely, tumour-core IL-17⁺ cell clustering was associated with higher cNLR without a clear OS association, whereas proximity of RORγ⁺IL-17⁺ARG1⁺ cells to CK7⁺ epithelium was associated with better survival without an evident relationship to NLR. Thus, tissue features associated with systemic inflammation were not necessarily those associated with clinical outcome. These relationships varied according to phenotype, anatomical location, and timing of the circulating measurement, arguing against a uniformly adverse interpretation of RORγ or IL-17 expression.

Building on prior evidence that RORC1 regulates tumour-associated EM (3, 13), our marrow findings provide temporal evidence supporting this mechanism. RORγ inverse agonism altered marrow progenitor composition, including a numerical reduction in eGMP frequency, before detectable divergence in tumour volume. This timing makes an explanation based solely on reduced tumour burden less likely, although it does not establish that eGMP changes mediate tumour control. At later endpoints, circulating neutrophil and RORγ⁺ neutrophil frequencies did not show clear treatment-associated reductions. These findings underscore a key distinction between dynamic myeloid programming and endpoint population measurements: circulating neutrophil fractions may remain relatively stable even when upstream hematopoiesis and tissue organization have already changed. The endpoint therefore does not necessarily capture the earlier marrow response or subsequent tumour trajectory.

The experimental metastasis studies further demonstrate that the consequences of RORγ perturbation depend on biological context. TF-S10 increased pulmonary metastatic burden after intravenous tumour-cell delivery in mice without prior tumour conditioning, whereas treatment initiated before intravenous challenge in mice bearing established flank tumours substantially reduced pulmonary burden. Delayed treatment produced less protection at the fixed endpoint. These contrasting outcomes support context-dependent effects of RORγ inhibition. They are compatible with an effect on host conditions encountered by disseminated tumour cells, although differences between the experimental schedules prevent attribution of the contrast exclusively to tumour conditioning.

The delayed-treatment findings also warrant a distinction between metastatic nodule number and overall pulmonary burden. An exploratory comparison identified lower lung weight with delayed treatment relative to controls, without a detectable difference in nodule count. The multiplicity-adjusted analysis did not identify a significant lung-weight difference between these groups. This observation leaves open the possibility that intervention after tumour-cell delivery affects pulmonary burden through changes not captured by nodule number alone. However, endpoint nodule counts and lung weights cannot establish altered growth velocity or distinguish effects on initial seeding from subsequent outgrowth. Differences in cumulative exposure further limit separation of treatment timing from treatment duration. The experiments therefore identify contrasting responses across conditioning and treatment schedules while leaving the underlying temporal mechanism unresolved.

Exploratory analyses examined tissue changes beyond those captured by circulating immune measurements. Across separately sampled murine tumour, marrow and histologically uninvolved lung cohorts, RORγ inverse agonism was associated with changes in local myeloid representation and organization, including reduced PD-L1 intensity in tumour-associated CD163⁺ myeloid cells and altered spatial relationships involving neutrophils, CD8⁺ T cells, and pulmonary macrophages. Together with the early marrow findings, these observations support tissue-specific consequences of perturbing myeloid regulation. Such effects may reflect changes in the differentiation, recruitment, and local interactions that generate and maintain cellular phenotypes. A phenotype observed at a single time point may therefore represent a dynamic programme whose function depends on anatomical and disease context; its association with outcome does not establish a persistently harmful function or imply that eliminating the expressing cells would be beneficial. Targeting upstream regulatory processes may instead reshape multiple cellular interactions over time without broadly depleting marker-defined populations. Our findings are consistent with this possibility, but systemic treatment and separate tissue cohorts cannot distinguish marrow-mediated from direct peripheral effects or establish coordinated responses within animals.

The tumour spatial findings prompted testing RORγ inverse agonism with PD-L1 blockade. TF-S10 produced more sustained tumour control than anti-PD-L1 alone across independent cohorts, consistent with prior work showing that RORγ inhibition can enhance checkpoint blockade in the setting of myeloid immunosuppression (13). These findings support further investigation of myeloid regulatory pathways as modifiers of the tissue environment in which checkpoint blockade operates. The mechanism underlying the combination benefit remains unresolved, however. Reduced myeloid PD-L1 intensity and altered neutrophil–CD8⁺ T-cell organization are plausible contributors, but the present data do not establish causality, and spatial-pair measurements must be interpreted alongside changes in cell abundance.

These findings inform interventions targeting the tumour–host response. The absence of substantial tumour-cell-autonomous activity in the assays performed supports a host contribution to TF-S10 activity, although direct tumour effects in vivo cannot be excluded. Importantly, the increased pulmonary burden observed in unconditioned mice demonstrates that molecular specificity alone does not predict therapeutic effect. The biological state of the host, tumour conditioning, disease phase, and treatment timing may all influence the consequences of perturbing myeloid regulation. cNLR provides an accessible clinical measure that could be evaluated alongside tumour exposure, treatment schedule, and tissue immune features, but these data do not establish cNLR as a predictor of treatment benefit or define a clinical treatment window.

Several limitations should be considered. The human cohorts were small and incompletely overlapping, limiting integrated patient-level analyses and multivariable adjustment. Survival analyses were exploratory and model discrimination was assessed within the same cohort used for model fitting. ARG1 served as an operational marker of the tissue neutrophil compartment but does not fully establish lineage or functional state. Murine spatial analyses included few biological replicates, and nested statistical models cannot overcome this limitation. Marrow and circulating frequencies also do not directly measure production rates or absolute cell numbers. Finally, the studies used syngeneic models and a single pharmacologic inverse agonist, and systemic treatment cannot assign effects exclusively to myeloid cells or resolve the contributions of RORγ versus RORγt. Intravenous challenge models likewise interrogate selected steps of pulmonary colonization rather than the complete metastatic cascade. Complementary pharmacological approaches, selective cellular perturbations and lineage studies are needed to clarify which components of the host response contribute to treatment efficacy.

In summary, RORγ-associated myeloid programming links systemic inflammation with hematopoiesis, tissue immune organization, and tumour progression in GEA. Its effects vary with anatomical context, tumour conditioning, and treatment timing, emphasizing that myeloid phenotypes should be interpreted as dynamic features of an evolving tumour–host system. These findings support identifying contexts where RORγ-associated regulation sustains adverse host responses and developing biomarkers integrating systemic, tissue and temporal features to guide treatment.

## METHODS

### 1.1 Human cohorts and biospecimens

Patients with histologically confirmed gastroesophageal adenocarcinoma (GEA) diagnosed between April 2014 and October 2018 were identified from the McGill University Health Centre Esophageal and Gastric Data and Bio-Bank. Patients with concurrent inflammatory or infectious conditions, another active malignancy at diagnosis, or squamous-cell carcinoma were excluded. Peripheral blood used for immune phenotyping was collected before systemic anticancer therapy. Circulating neutrophil-to-lymphocyte ratio (cNLR) was calculated from contemporaneous absolute neutrophil and lymphocyte counts and classified as low (<4) or high (≥4) using a prespecified threshold(23–25).

Tissue analyses used an established 39-patient spatial TMA cohort, of which 29 patients had evaluable tissue for the RORγ-focused analyses. TMA cores were assigned to tumour core, tumour periphery, normal stomach and normal esophagus. Methods detail and clinicopathological characteristics are provided in **Supplementary Methods S1 and Table S1**.

The Esophageal and Gastric Data and Bio-Bank is registered under protocol MP-37-2007-856, and associated experimental studies were approved by the McGill University Health Centre Research Ethics Board under protocol 2024-9907. All participants provided written informed consent.

### 1.2 Human RORγ phenotyping

#### 1.2.1 Circulating neutrophil flow cytometry

Treatment-naïve peripheral blood was analysed from patients with low (n=18) or high (n=8) cNLR. Following erythrocyte lysis, viable CD66b⁺CD16⁺ neutrophils were identified by flow cytometry and assessed for intracellular RORγ and IL-17A and surface PD-L1 expression. RORγ⁺ and RORγ⁻ populations were defined using fluorescence-minus-one and appropriate isotype controls. Samples were acquired on a BD LSRFortessa and analysed using FlowJo v10.10 (BD Biosciences). Antibodies, clones, fluorophores and staining conditions are provided in **Supplementary Methods S1.1**.

#### 1.2.2 Multiplex tissue immunofluorescence and image analysis

Formalin-fixed paraffin-embedded TMA sections were stained with Akoya Biosciences® Opal™ multiplex automation kit and were imaged on Vectra® 3.0 multispectral imaging system (Akoya/PerkinElmer). Cellular segmentation and phenotyping were performed in HALO v4. (Indica Labs) using DAPI-based segmentation and prespecified consistent marker thresholds. ARG1⁺ myeloid cells were subsequently classified according to RORγ and IL-17 expression. Analyses were performed within predefined anatomical compartments and summarized at the patient level.

Detailed staining and image-analysis procedures are provided in **Supplementary Methods S1.2 and Supplementary Table S2.**

#### 1.2.3 Human spatial and survival analyses

Core-level abundance, composition and spatial measurements were summarized as patient–compartment medians. Spatial endpoints included occupied-tile fraction, tile-prevalence variability, RORγ⁺ARG1⁺ enrichment within ARG1-rich hotspots, IL-17⁺ spatial autocorrelation and proximity to CK7⁺ epithelial cells. Clinical associations used patient-level observations, with Spearman correlation for cNLR and continuous Cox models for survival. The tumour-periphery enrichment model used a postoperative day-180 landmark, five-year postoperative censoring and adjustment for matched log-transformed cell abundance. Median-split Kaplan–Meier curves provided illustrative displays. Human spatial definitions, endpoint-specific eligibility, clinical sampling windows, correction families and survival models are provided in **Supplementary Methods S1.3–S1.7**.

### 1.3 Animals, tumour cell lines and RORγ inverse agonist

Male and female C57BL/6 mice aged 7–10 weeks (Charles River Laboratories) were used. Procedures followed institutional guidelines and were approved by the RI-MUHC Animal Research Division (protocol 8210). Syngeneic models used YTN16-GFP gastric, MC38-RFP colorectal and H59-GFP lung cancer cells, provided by Drs. Goldenring, N. Beauchemin and P. Brodt, respectively. Culture conditions are detailed in Supplementary Methods S2.

TF-S10 is a synthetic small-molecule inverse agonist targeting the shared RORγ ligand-binding domain, provided by Dr Tchervenkov’s Transplant Immunology Laboratory (26). TF-S10 or vehicle was administered subcutaneously once daily at the model-specific doses and schedules described below and in the figure legends.

### 1.4 Preclinical tumour models and RORγ perturbation

#### 1.4.1 Established YTN16 gastric tumour model

Mice received 5 × 10⁶ YTN16-GFP cells subcutaneously in the flank. TF-S10 or vehicle was initiated after tumour establishment on D7, and tumour growth was followed longitudinally. At endpoint (D38), tumour, blood and lung were collected for immune phenotyping and assessment of spontaneous pulmonary dissemination.

#### 1.4.2 MC38 primary tumour and temporal myelopoiesis model

Mice received 2.5 × 10⁵ MC38-RFP cells subcutaneously, with TF-S10 or vehicle initiated on D7. Complementary timed-collection and D28 endpoint cohorts were used to relate tumour progression to circulating myeloid populations and bone-marrow progenitor composition.

Mice received intraperitoneal anti-PD-L1 every three days from D7 until endpoint; combination-treated mice also received daily TF-S10.

#### 1.4.3 Tumour-conditioned MC38 metastasis model

Mice bearing subcutaneous MC38-RFP tumours received 2.5 × 10⁵ MC38-RFP cells intravenously on D14. T TF-S10 began before (D7) or after (D17) metastatic challenge, with vehicle-treated controls. Pulmonary metastatic burden, blood and bone-marrow populations were analysed at endpoint.

#### 1.4.4 Unconditioned MC38 tail-vein metastasis model

Tumour-free mice received 2.5 × 10⁵ MC38-RFP cells intravenously followed by TF-S10 or vehicle beginning 24 hours later. Pulmonary metastatic burden was assessed at D28 together with circulating and marrow immune populations.

#### 1.4.5 Cross-compartment spatial model

Tumour, BM and histologically uninvolved lung from TF-S10- and vehicle-treated MC38-RFP bearing mice were analysed by immunofluorescence and spatial profiling at D28. Multiple regions were sampled within each tissue, with the mouse or specimen retained as the biological replicate. Detailed image acquisition, cell classification and spatial-analysis procedures are described in **section 1.7**.

Detailed experimental procedures, including treatment dosing, sample collection, tissue processing and model-specific endpoint assessments, are provided in the **Supplementary Methods S3.**

### 1.5 In-vitro characterization of tumour-cell RORγ expression and TF-S10 response

RORγ expression in YTN16-GFP, MC38-RFP and H59-GFP tumour cells was assessed using immunofluorescence, intracellular flow cytometry and/or immunoblotting, with RORγ-expressing cells (Hep55.1c) or tissues (thymus) included as positive controls. Direct tumour-cell responses to TF-S10 were assessed using clonogenic growth, metabolic viability, scratch-closure and Annexin V/propidium iodide apoptosis assays.

Detailed protocols are indicated in **Supplementary Methods S4 and Supplementary Fig. S2-3**.

### 1.6 Murine sample processing and flow cytometry

Peripheral blood underwent erythrocyte lysis followed by multicolour flow cytometry to quantify circulating Ly6G⁺CD11b⁺ neutrophils and RORγ-associated immune phenotypes. Bone-marrow from femora and tibiae was processed into single-cell suspensions and analysed using a lineage and progenitor panel. Conventional myeloid progenitors were identified within the lineage-negative Sca-1⁻CD117⁺ compartment, whereas eGMPs were defined within lineage-negative Sca-1⁺CD117⁺ cells as shown in **Figure 4**.

Samples were acquired using a BD LSRFortessa and analysed in FlowJo v10.10 (BD Biosciences). Complete antibody panels, clones, dilutions, gating controls and processing procedures are provided in **Supplementary Methods S5.**

### 1.7 Tissue immunofluorescence and spatial analysis

Tumour, lung and bone-marrow sections were stained using experiment-specific immunofluorescence or immunohistochemical panels. Images were analysed using HALO, inForm or ImageJ, with QuPath used for registration checks and visualization.

For histologically uninvolved lung (Fig. 6G–H), HALO v4.2.2 provided phenotype calls, Ly6G nuclear optical-density (OD) values and coordinates. Review of 144 coordinate-matched candidate neutrophil objects against source images informed a fixed criterion requiring both a positive HALO neutrophil call and Ly6G nuclear OD ≥0.130. Review was blinded to mouse identity, treatment assignment, OD values and treatment results. All 144 objects in a separate blinded verification sample (24 per mouse) passed review for Ly6G signal and compatible morphology. The criterion was applied uniformly without modifying original HALO calls, measurements or coordinates; macrophages retained their HALO classification.

Spatial analyses used systematic sampling within each tissue, with fields or tiles treated as subsamples and mice or marrow specimens as biological replicates. Primary comparisons used replicate-level summaries; supporting mixed-effects or cluster-aware analyses accounted for within-replicate dependence (27–29). Staining panels, processing workflows, eligibility thresholds, grid dimensions, spatial metrics and sensitivity analyses are detailed in **Supplementary Methods S6–S7** and **Supplementary Tables S3–S5.**

### 1.8 Statistical analysis

Statistical analyses used GraphPad Prism v10.6.1 and R v4.5.1. Tests were two-sided with *P*<0.05 considered statistically significant. Two-group comparisons used paired or unpaired parametric/non-parametric tests as appropriate, and continuous associations were assessed using Spearman correlation. Survival analyses used Kaplan–Meier estimates and Cox proportional-hazards regression.

Longitudinal analyses used treatment-by-time models where indicated. For spatial analyses, measurements nested within the same mouse or specimen were analysed using replicate-level summaries or models accounting for within-replicate dependence. Exact tests and sample sizes are reported in the corresponding figure legends.

## DECLARATIONS

### Ethics approval and consent to participate

The Esophageal and Gastric Data- and Biobank is registered under Nagano protocol MP-37-2007-856. The biobank and all associated translational research studies were approved by McGill University Health Centre Research Ethics Board (REB protocol 2024-9907). Written informed consent was obtained from all participants. Mouse experiments were approved by the Research Institute of McGill University Health Centre Animal Research Division under protocol 8210.

### Data availability

All data supporting the findings of this study are available from the corresponding author upon reasonable request.

## Acknowledgements

We acknowledge the patients who participated in this study and the Thoracic Surgery team at the Montreal General Hospital for their support with patient recruitment, consenting and biospecimen collection. We also acknowledge the Research Institute of McGill University Health Centre for providing the infrastructure and resources that enabled this research, the RI-MUHC Molecular Imaging Core Platform, the RI-MUHC Immunophenotyping platform and RI-MUHC Animal Research Division for their expertise. Further acknowledgements go to the members of the Experimental Pathology Research Laboratory [RRID:SCR_017928] at NYU Langone’s Laura and Isaac Perlmutter Cancer Center for their assistance with staining and multispectral imaging of the human tissue microarrays. The Experimental Pathology Research Laboratory is partially supported by the Cancer Center Support Grant P30CA016087. The PhenoImagerHT multispectral imaging system used in this study was acquired through Shared Instrumentation Grant S10 OD021747.

