## Supplemental Material for "RORγ Inverse Agonism Redirects Tumour-Conditioned Emergency Myelopoiesis and Attenuates Established Tumour Progression"

### SUPPLEMENTAL METHODS

#### S1. Human cohorts, tissue classification and biospecimen processing

Peripheral blood specimens used for flow-cytometric analyses were obtained before initiation of systemic anticancer therapy. Peripheral blood was collected into EDTA-containing tubes and processed for downstream immune phenotyping as described below.

Human tissue analyses used an established 39-patient multiplex spatial tissue microarray (TMA) cohort. Endoscopic biopsy or surgical resection specimens containing tumour and, where available, adjacent non-malignant tissue were reviewed histologically before TMA construction, with tissue regions selected by a pathologist from hematoxylin and eosin-stained sections. The present RORγ-focused study used separate tissue sections and study-specific staining, cellular phenotyping and spatial analyses.

Individual TMA cores were linked to patients and anatomical compartments using the locked TMA core map and histopathological classifications. Compartments included tumour core, defined as central malignant tissue; tumour periphery, representing the tumour–stroma interface; histologically normal stomach and oesophagus without tumour involvement; Barrett's oesophagus or dysplasia; and lymph-node metastases. Analyses in the main study were restricted to the compartments relevant to each prespecified comparison.

Formalin-fixed, paraffin-embedded TMA specimens were analysed retrospectively using de-identified clinical and pathological annotations. All biospecimens were de-identified before experimental analysis.

##### S1.1 Peripheral-blood flow cytometry

Peripheral blood from treatment-naïve patients with GEA was collected into EDTA-containing tubes. Whole blood underwent erythrocyte lysis using 1× BD Pharm Lyse Lysing Buffer (Cat. 555899) for 10 min at room temperature in the dark, followed by centrifugation at 450 × *g* for 5 min. The resulting leukocyte pellet was stained with fixable eBioscience eFluor 780 viability dye (Cat. 65-0865-14; 1:1000) for 10 min on ice.

Cells were stained for the surface markers CD66b-APC (Miltenyi Biotec; clone REA306; Cat. 130-135-357), CD16-VioGreen (Miltenyi Biotec; clone REA423; Cat. 130-

113-959) and PD-L1-PE-Cy7 (eBioscience; clone M1H1; Cat. 25-5983-41) for 30 min on ice in the dark.

For intracellular staining, cells were fixed and permeabilized using the eBioscience FoxP3/Transcription Factor Staining Buffer Set (Cat. 00-5523-00) according to the manufacturer's instructions and stained with RORγ-PE (eBioscience; clone AFKJS-9; Cat. 12-6988-82; 1:50) and IL-17A-BV711 (BioLegend; clone BL168; Cat. 512327; 1:20) for 30 min at room temperature.

Single-stained compensation controls were prepared using antibody-capture beads or individually stained cells. Fluorescence-minus-one controls were used to establish PD-L1 gates. For RORγ and IL-17A, the corresponding fluorescence-minus-one controls included matched isotype controls and were used to define positivity thresholds.

Samples were acquired on a BD LSRFortessa flow cytometer and analysed using FlowJo v10.10 (BD Biosciences). Neutrophils were identified as viable $CD66b^+CD16^+$ cells, with $ROR\gamma^+$ and $ROR\gamma^-$ populations subsequently assessed for IL-17A and PD-L1 expression according to the gating strategy shown in **Figure 1**.

#### S1.2 Multiplex tissue immunofluorescence and image analysis

Formalin-fixed, paraffin-embedded TMA sections of five μm each were immunostained on a Leica BondRx auto-stainer with Akoya Biosciences® Opal™ multiplex automation kit reagents according to the manufacturer's instructions. In brief, sections were deparaffiinized online and treated with 3% $H_2O_2$ (Thermo Scientific, Cat: 426000010) to inhibit endogenous peroxidases, followed by antigen retrieval with either ER1 (Leica, Cat AR9961; pH6) or ER2 (Leica, Cat: AR9640; pH9) retrieval buffer at 100C for 20 minutes. After blocking with Primary Antibody Diluent (Leica, Cat: AR93520), slides were incubated with the first primary antibody and secondary HRP polymer pair, followed by HRP-mediated tyramide signal amplification with a specific Opal® fluorophore. Once the Opal® fluorophore was covalently linked to the antigen, primary and secondary antibodies were removed with a heat retrieval step. This sequence was repeated 5 more times with subsequent primary and secondary antibody pairs, using a different Opal fluorophore with each primary antibody (**Supplementary Table S2**). After antibody staining, sections were counterstained with spectral DAPI (Akoya Biosciences, Cat: FP1490) and mounted with ProLong Gold Antifade (Thermo Scientific, Cat: P36935).

Semi-automated image acquisition was performed on a Vectra® 3.0 multispectral imaging system (Akoya/PerkinElmer). After an initial whole slide scan, tissues were manually outlined to select fields for multispectral imaging at 20X. The fluorophore emission signatures were captured by a multispectral camera and then unmixed with InForm® software version 2.4.2 (Akoya/PerkinElmer). Autofluorescence was removed from the composite images and pseudo-colored images were exported as .tif files using InForm®'s proprietary algorithm.

Image processing and quantitative analysis were performed in HALO (Indica Labs) using the Multiplex IF module. Spectral unmixing was used to separate individual fluorophore signals and generate single-marker and composite images. Cellular segmentation was based on DAPI nuclear staining. Marker-positivity thresholds were manually optimized using representative positive and negative regions and then applied uniformly across all samples within the staining cohort.

$ARG1^{+}$ cells were used to operationally define the myeloid population of interest, with RORγ and IL-17 expression subsequently used to resolve type-17-associated phenotypes. Cellular phenotypes, marker co-expression, cell counts and spatial coordinates were quantified within predefined anatomical compartments, including tumour core, tumour periphery, normal stomach and normal oesophagus.

Spatial analyses were performed using the patient-linked anatomical assignment of each TMA core. Where multiple cores contributed to the same patient and compartment, measurements were summarized at the patient level for paired tissue comparisons and associations with clinical variables and outcome. Representative images were selected from the same quantified datasets and were not treated as independent observations.

Human spatial metrics, patient-level aggregation, clinical associations and survival analyses are described in Supplementary Methods S1.3–S1.7.

#### S1.3 Human spatial measurements and patient-level aggregation

Cellular phenotypes and X–Y positions were linked to the patient and anatomical compartment assigned to each TMA core. Exported cell positions were calculated as the midpoints of the object bounding boxes. Core-level measurements were summarized using the median across evaluable cores within each patient and compartment. The patient was the unit of inference; cores and spatial tiles were subsamples. $ARG1^+$ abundance was expressed as cell count per core. $ROR\gamma^+$ composition was calculated as the number of $ROR\gamma^+ARG1^+$ cells divided by the number of $ARG1^+$ cells in each core, excluding zero denominators, before patient-level aggregation.

For the normal-stomach versus tumour-periphery coverage and patchiness comparison, tissue was partitioned into square tiles with a width of 250 native coordinate units. The grid was anchored to the minimum object XMin and YMin coordinates within each core, and cells were assigned to tiles according to their bounding-box midpoints. Native coordinates were retained for computation. All three slides used a calibration of approximately 0.496 μm per pixel, corresponding to a tile width of approximately 124 μm.

Eligible tiles contained at least 20 segmented cells. Boundary tiles were retained if they met the same cell-count criterion; no explicit tissue mask or correction for partial tile area was applied. Eligible cores contained at least four eligible tiles and at least five phenotype-positive cells summed across those tiles. Tile prevalence was calculated as the number of phenotype-positive cells divided by the total number of segmented cells within each eligible tile. Occupied-tile fraction was the proportion of eligible tiles containing at least one phenotype-positive cell. Patchiness was summarized as the sample standard deviation of tile prevalence divided by its mean, including eligible tiles with zero phenotype-positive cells. Values with zero mean prevalence were treated as undefined and recorded as missing.

Core-level spatial measurements were aggregated to patient–compartment medians, and paired comparisons included patients with evaluable measurements in both compartments. Comparisons used two-sided paired Wilcoxon signed-rank tests, with Benjamini–Hochberg correction applied separately within each phenotype–metric family across eight tissue comparisons: normal stomach versus normal oesophagus; tumour core versus normal oesophagus; tumour periphery versus normal oesophagus; tumour core versus normal stomach; tumour periphery versus normal stomach; tumour periphery versus tumour core; lymph-node metastasis versus tumour core; and lymph-node metastasis versus tumour periphery. Sensitivity analyses repeated the spatial calculations using tile widths of 200 and 500 native coordinate units (approximately 99.2 and 248 μm, respectively), retaining the same eligibility and aggregation rules and examining changes in compartment-level summaries and evaluability.

#### S1.4 RORγ⁺ARG1⁺ enrichment within ARG1-rich hotspots

Hotspot enrichment was calculated using a separate square grid with bin widths of 50 native coordinate units in the original HALO export. The grid was anchored to the minimum X and Y bounding-box midpoint coordinates across all segmented cells within each core. Native coordinates were retained for computation. All three slides used a calibration of approximately 0.496 µm per pixel, corresponding to a bin width of approximately 24.8 µm.

Bins containing $ARG1^+$ cells were ranked by $ARG1^+$ cell count. The number of hotspot bins was defined as 20% of the ARG1-positive bins, rounded upward, and the highest-ranking bins were selected; ties were resolved by bin identifier. Core eligibility required at least five $ARG1^+$ cells, at least five occupied bins across all input cells and at least three bins containing $ARG1^+$ cells.

For each eligible core, enrichment was calculated as $\log_2\{[(R\_in + 0.5)/(A\_in + 1)]/[(R\_out + 0.5)/(A\_out + 1)]\}$, where R and A denote $ROR\gamma^+ARG1^+$ and $ARG1^+$ cell counts, respectively, inside or outside the selected hotspots. The pseudocounts stabilized estimates when counts were zero. Positive enrichment values indicated a higher $ROR\gamma^+$ fraction among $ARG1^+$ cells inside ARG1-rich hotspots than outside them. Core-level enrichment values were summarized as medians across evaluable cores within each patient and anatomical compartment.

#### S1.5 Spatial autocorrelation and epithelial proximity

Tumour-core $IL\text{-}17^+ARG1^+$ spatial organization was assessed using global Moran's I, calculated from phenotype prevalence across square tiles of 500 native coordinate units using the grid definition and eligibility criteria described in S1.3. Each eligible tile contained at least 20 segmented cells, and evaluable cores contained at least four eligible tiles and five phenotype-positive cells summed across those tiles. Tile prevalence was defined as the number of phenotype-positive cells divided by the total number of segmented cells within the tile.

Moran's I was calculated as:

$$I = \frac{n}{S_0} \frac{\sum_i \sum_j w_{ij}(p_i - \bar{p})(p_j - \bar{p})}{\sum_i (p_i - \bar{p})^2}, \qquad S_0 = \sum_i \sum_j w_{ij}.$$

where (n) denotes the number of eligible tiles, (p_i) the phenotype prevalence in tile (i), and (\bar p) the mean prevalence across eligible tiles. Binary queen-adjacency weights assigned (w_{ij}=1) to distinct eligible tiles sharing an edge or corner and zero otherwise, with (w_{ii}=0). Weights were not row-standardized; normalization was provided by the (n/S_0) factor. Disconnected components were not joined by additional spatial links. Isolated eligible tiles remained in the tile count, mean and variance calculations but contributed no neighbouring pairs to the numerator. Regions with zero phenotype prevalence had undefined Moran's I values and were recorded as missing. The statistic is also undefined when tile prevalence has zero variance or no neighbouring tile pairs are available. Core-level values were summarized as patient–compartment medians.

Epithelial proximity was calculated from the Euclidean distance between each RORγ$^+$IL-17$^+$ARG1$^+$ cell's bounding-box midpoint and the midpoint of its nearest operational CK7$^+$ epithelial cell within the same core. CK7 positivity was defined by an Opal 780 Positive Classification value equal to 1. For each distance threshold, the numerator was the number of target cells whose nearest CK7$^+$ cell lay at or within that threshold, and the denominator was the total number of RORγ$^+$IL-17$^+$ARG1$^+$ cells in the core. Each target cell contributed once, irrespective of the number of nearby CK7$^+$ cells.

Primary proximity analyses required at least five target cells and one CK7$^+$ cell per core. Cores without CK7$^+$ cells or with insufficient target cells were excluded rather than assigned zero proximity. A sensitivity analysis required at least 20 CK7$^+$ cells while retaining the five-target-cell minimum. Core-level proximity proportions were summarized as patient–compartment medians for clinical associations.

The primary proximity threshold was 20 native coordinate units, with corresponding 50- and 100-unit measurements included in the focused survival-analysis family. These distances were calculated in the original coordinate system. At the verified LF-EAC4 calibration, the thresholds corresponded to approximately 9.92, 24.80 and 49.60 μm, respectively; this calibration was not assumed to apply to other slides.

### S1.6 Clinical sampling windows, comparisons and multiple testing

Preoperative cNLR was obtained from the CBC closest to surgery within the preceding 30 days, excluding the day of surgery. Recovery-window cNLR was the patient median during postoperative days 60–180 inclusive, displayed as approximately 0.16–0.49 years. Associations between continuous tissue features and cNLR were assessed using two-sided Spearman correlation. The cNLR threshold of 4 was used for illustrative grouping in the reported correlation plots; correlation tests retained continuous values.

Paired ARG1$^{+}$ abundance comparisons used two-sided Wilcoxon signed-rank tests with Benjamini–Hochberg correction across the original six-comparison family: normal stomach versus normal oesophagus, tumour core versus normal stomach, tumour periphery versus normal stomach, tumour periphery versus tumour core, lymph-node metastasis versus tumour core, and lymph-node metastasis versus tumour periphery. The paired normal-stomach versus tumour-periphery RORγ$^{+}$ composition comparison used a two-sided Wilcoxon signed-rank test.

For preoperative spatial–cNLR associations, Benjamini–Hochberg adjustment was recalculated across a revised exploratory family of 16 features, defined retrospectively during the reproducibility review and documented in the accompanying package. For recovery-window cNLR, the reported IL-17$^{+}$ Moran's I association used Benjamini–Hochberg correction across 11 tests: measurements in normal oesophagus, normal stomach, tumour core, tumour periphery and lymph-node metastasis, together with six within-patient differences—normal stomach minus normal oesophagus, tumour core minus normal stomach, tumour periphery minus normal stomach, tumour periphery minus tumour core, lymph-node metastasis minus tumour core, and lymph-node metastasis minus tumour periphery. This family yielded $q = 0.002568$ for tumour-core IL-17$^{+}$ Moran's I. The separate six-feature recovery-window analysis yielded $q = 0.0014007$ and did not supply the adjusted value reported in the final figure.

The recovery-window correction followed the original analysis structure, grouping tests within each cNLR exposure and spatial domain across anatomical compartments and contrasts. The original correction families were retained when displaying selected associations, so multiplicity adjustment was not restricted to the panels shown.

Analyses used available patient-level observations for each endpoint, without imputation. Correlations required evaluable tissue measurements and cNLR within the relevant sampling window; paired comparisons and anatomical differences additionally required evaluable measurements in both compartments. Consequently, sample sizes varied between tests and were reported for each association. Missing or undefined spatial measurements were excluded from the relevant analysis rather than assigned a value of zero.

Clinical association analyses were exploratory. In the original recovery-window analysis, associations with at least eight evaluable patients were considered potentially interpretable exploratory findings, those with five to seven patients were treated as sensitivity analyses, and those with four or fewer patients were descriptive only. The small-sample tests remained within the original multiplicity family. The available analysis records do not establish prospective prespecification of the displayed associations.

### S1.7 Human survival analyses

Survival analyses were performed in R version 4.5.1 using the survival package version 3.8.9. Continuous spatial measurements were evaluated using Cox regression with Efron handling of tied event times. Hazard ratios and 95% confidence intervals were reported per one-standard-deviation increase. Analyses used complete patient-level observations without imputation. Median-split Kaplan–Meier curves and log-rank tests provided illustrative comparisons distinct from the continuous-model estimates.

The tumour-periphery hotspot-enrichment analysis used a postoperative day-180 landmark and included patients alive and followed beyond day 180 with both enrichment and matched abundance available. Follow-up was censored at five years after surgery, using 365.25 days per year. Analysis time was min(postoperative survival days, 1826.25) − 180; deaths after five years were treated as censored. Hotspot enrichment and $\log_2(1 + \text{ROR}\gamma^+\text{ARG1}^+$ tumour-periphery cell count) were standardized using the cohort mean and sample standard deviation and entered together into a Cox model. A one-degree-of-freedom likelihood-ratio test compared this model with the abundance-only model. The analysis included 23 patients and 10 deaths.

Epithelial-proximity analyses used the supplied overall-survival duration, capped at five years, without a postoperative landmark. Separate univariable Cox models retained the original endpoint-specific standardization. Eligibility required at least eight patients, three deaths, both events and censored observations, and non-zero biomarker variation. Benjamini–Hochberg correction was applied across four endpoints: tumour-core RORγ$^{+}$IL-17$^{+}$ARG1$^{+}$ proximity to CK7$^{+}$ cells at thresholds of approximately 9.92, 24.80 and 49.60 μm (20, 50 and 100 native coordinate units, respectively), and tumour-periphery occupied-tile fraction.

Tumour-core IL-17$^{+}$ Moran's I was evaluated from surgery through five years in 24 patients with 12 deaths. An exploratory post hoc analysis divided follow-up at two years, retaining the full-cohort biomarker standardization. All 24 patients contributed to the first interval; 18 patients alive and followed beyond two years entered the second risk set. The models contained no additional covariates. Hazard ratios were 1.88 (95% CI, 0.99–3.57) during years 0–2 and 0.46 (95% CI, 0.16–1.28) during years 2–5, with six deaths in each interval. A one-degree-of-freedom likelihood-ratio test compared interval-specific coefficients with a common coefficient; the interaction P value was nominal.

Sensitivity analyses for the original full-follow-up proximity models included interquartile-range scaling, leave-one-patient-out refitting with restandardization, and exploratory adjustment for ordinal stage I–IV, prioritizing pathological over clinical stage. An endpoint-by-log-time interaction provided a proportional-hazards sensitivity check. These analyses were separate from the final five-year refits.

Harrell's C-index was obtained from the Cox-model concordance output. Five-year discrimination compared predicted risk between patients who died within five years and those observed alive through five years; patients censored earlier had unknown five-year status. The C-index and five-year AUC were interpreted as apparent performance in the model-fitting cohort, without independent validation.

### S2. Animals, tumour cell lines and RORγ inverse agonist

#### S2.1 Animals

C57BL/6 mice (Charles River Laboratories), aged 7–10 weeks, were used for all in vivo experiments. Male and female mice were included in experimental cohorts. Animals were maintained under standard housing conditions with access to food and water *ad libitum*. Experimental procedures were performed under protocol 8210 in accordance with the Veterinary Authority of the Institutional Animal Care and Use Committee at the Research Institute of the McGill University Health Centre (RI-MUHC).

#### S2.2 Murine tumour cell lines and culture

YTN16-GFP (Dr Goldenring, Vanderbilt University Medical Centre, USA), MC38-RFP (Dr N. Beauchemin, McGill University, Canada) and H59-GFP (Dr P. Brodt, RI-MUHC, Canada) murine tumour cell lines were used as syngeneic models of gastric, colorectal and lung cancer, respectively.

YTN16-GFP cells were maintained in DMEM containing 4.5 g/L glucose, L-glutamine and sodium pyruvate (Wisent; Cat. 319-005-CL), supplemented with 10% fetal bovine serum (Wisent; Cat. 090150), 0.5% penicillin/streptomycin (Wisent; Cat. 450-201-EL) and 0.1% MITO+ serum extender (Corning; Cat. 355006). Cells were cultured on rat-tail collagen I-coated cultureware (Sigma-Aldrich; Cat. C3867).

MC38-RFP and H59-GFP cells were maintained in DMEM containing 4.5 g/L glucose, L-glutamine and sodium pyruvate (Wisent; Cat. 319-005-CL), supplemented with 10% fetal bovine serum and 1% penicillin/streptomycin. Cells were maintained at 37°C in a humidified atmosphere containing 5% $CO_2$.

Hep55.1c cells were used only as a positive control for RORγ expression in the in-vitro characterization experiments.

#### S2.3 RORγ inverse agonist

TF-S10 (Transplant Immunology Laboratory of Dr Tchervenkov, RI-MUHC) is a synthetic small-molecule RORγ inverse agonist belonging to a series of 2,3-disubstituted 4,5,6,7-tetrahydrobenzothiophene derivatives (molecular weight, 426.57 g/mol) and was used for pharmacological perturbation of RORγ-associated transcription [38]. TF-S10 was originally characterized using RORγt ligand-binding and type-17 functional assays. Because RORγ1 and RORγt share the ligand-binding domain targeted by pharmacological RORγ modulators, TF-S10 effects in the present study were interpreted as RORγ-associated pharmacological perturbation rather than isoform-specific RORγt inhibition.

For in vivo studies, TF-S10 was administered by subcutaneous injection once daily. Unless otherwise specified for an individual experimental model, mice received TF-S10 at 25 mg/kg, while control animals received the corresponding vehicle consisting of PBS containing 4% Tween 80 (Sigma-Aldrich, cat P1754). Treatment initiation and duration varied according to the individual experimental model and are described in the corresponding model-specific Supplementary Methods and figure legends.

### S3. Preclinical tumour models

#### S3.1 Established YTN16 gastric tumour model

C57BL/6 mice were inoculated subcutaneously in the left flank with 5 × $10^6$ YTN16-GFP cells. Treatment was initiated on day 7 after tumour inoculation, once tumours were established. Mice received daily subcutaneous TF-S10 or the corresponding vehicle according to the dosing schedule specified for the experiment. Tumour dimensions were measured longitudinally until the predefined experimental endpoint.

At endpoint, peripheral blood, primary tumour and lungs were collected. Circulating and intratumoural myeloid populations were assessed by flow cytometry, and pulmonary tumour burden was evaluated using GFP-based imaging together with lung immune phenotyping. Serial blood samples were analysed where available to assess longitudinal changes in circulating neutrophil populations and cNLR.

#### S3.2 MC38-RFP primary tumour and temporal myelopoiesis model

C57BL/6 mice were inoculated subcutaneously in the left flank with 2.5 × $10^5$ MC38-RFP cells. TF-S10 treatment was initiated on day 7 after tumour inoculation and administered once daily; control mice received vehicle on the same schedule. Tumour volumes were measured longitudinally.

Complementary experimental cohorts were used to define the temporal relationship between RORγ perturbation, myelopoiesis and tumour progression. Timed-collection cohorts underwent terminal sampling at defined intermediate time points during tumour progression, while a fixed-endpoint cohort was followed to day 28. Peripheral blood was analysed for circulating $Ly6G^+CD11b^+$ and $ROR\gamma^+$ neutrophils, and bone marrow was collected from femora and tibiae for progenitor phenotyping.

Conventional myeloid progenitors and emergency granulocyte–monocyte progenitors (eGMPs) were quantified by flow cytometry. eGMP trajectories were compared with tumour-volume trajectories to determine whether treatment-associated changes in marrow myelopoiesis preceded, coincided with or followed divergence in tumour growth.

For anti-PDL1 monotherapy, mice were treated with 10mg/kg of InVivoPlus anti-mouse PD-L1 antibody (BioXCell; Cat: BP0101-100MG-A), administered intraperitoneally every three days from day 7 till endpoint. Corresponding controls received InVivoMAb rat IgG2b isotype control (BioXCell; Cat: BE0090-100MG-A) intraperitoneally on the same schedule. Antibodies were diluted to the required concentration (10mg/kg) using InVivoPure pH6.5 Dilution Buffer (BioXCell; Cat: IP0065-50ML-A).

Mice receiving combination therapy were treated with both 25mg/kg TF-S10 daily and 10mg/kg anti-PD-L1 every three days. Combination control mice received both RORγ vehicle control (4% tween80 + PBS) daily and IgG2b isotype control every three days. Tumour volumes were measured longitudinally.

#### S3.3 Tumour-conditioned MC38-RFP metastasis model

To model metastatic establishment in a tumour-conditioned host, C57BL/6 mice were inoculated subcutaneously with $2.5 \times 10^5$ MC38-RFP cells in the left flank. On day 14 after primary tumour inoculation, mice received $2.5 \times 10^5$ MC38-RFP cells intravenously by tail-vein injection to induce pulmonary metastatic seeding.

Mice were assigned to control, early-treatment or delayed-treatment groups. Early TF-S10 treatment was initiated on day 7, before intravenous tumour-cell challenge, whereas delayed treatment was initiated on day 17, after metastatic seeding. Treatment continued until endpoint according to the dosing schedule specified for the experiment.

Peripheral blood and bone marrow were collected for immune and progenitor analyses. Lungs were harvested at endpoint, and pulmonary metastatic burden was quantified by fluorescence imaging of RFP-positive lesions together with lung-weight and metastatic-nodule assessments where applicable.

#### S3.4 Unconditioned MC38-RFP tail-vein metastasis model

To assess metastatic establishment in the absence of prior primary-tumour conditioning, C57BL/6 mice received $2.5 \times 10^5$ MC38-RFP cells intravenously by tail-vein injection without preceding subcutaneous tumour implantation. TF-S10 or vehicle was initiated 24 hours after tumour-cell injection and continued until the day-28 endpoint.

At endpoint, lungs were collected and pulmonary metastatic burden was quantified by enumeration of visible metastatic nodules. Peripheral blood and bone marrow were also collected for flow-cytometric analysis of circulating immune populations and marrow myelopoiesis.

#### S3.5 Cross-compartment spatial model

A dedicated MC38-bearing cohort was used to evaluate spatial effects of RORγ perturbation across primary tumour, bone marrow and histologically uninvolved lung. Mice were inoculated subcutaneously with MC38-RFP cells and treated with TF-S10 or vehicle according to the established-treatment schedule.

At day 28, primary tumour, femoral/tibial bone marrow and lungs without histologically evident tumour involvement were collected for immunofluorescence and spatial analysis. Multiple microscopic regions or spatial tiles were sampled systematically from each tissue. These regions were treated as within-replicate spatial observations rather than independent biological replicates; the individual mouse, or bone-marrow specimen where applicable, remained the unit of biological replication.

Detailed tissue processing, cellular phenotyping, image acquisition, spatial sampling, metric definitions and nested statistical analyses are described in the **Supplementary Methods S6-7 and Supplementary Tables S3-5.**

### S4. Tumour-cell RORγ expression and direct TF-S10 response

#### S4.1 Tumour-cell RORγ expression

RORγ expression in murine tumour cell lines was assessed using complementary immunofluorescence, intracellular flow cytometry and immunoblotting approaches.

Hep55.1c cells and mouse thymus lysate were used as positive controls where appropriate.

For immunofluorescence, YTN16-GFP cells were seeded at 2 × $10^5$ cells per well in chamber slides and cultured to confluence. Cells were fixed in 4% paraformaldehyde overnight at 4°C, permeabilized with 0.1% Triton X-100 and blocked for 1 h in TBST containing 1% bovine serum albumin, 1% fetal bovine serum and 0.01% Triton X-100. Cells were stained with anti-mouse RORγ-PE (clone AFKJS-9; Cat. 12-6988-82; 1:50) together with DAPI, mounted using Vectashield Vibrance antifade medium and imaged using a Zeiss LSM 780 confocal microscope. Images were analysed using ImageJv2.1.

For immunoblotting, whole-cell lysates from YTN16-GFP cells were prepared in RIPA buffer (Thermo Scientific; Cat. 89900) supplemented with protease and phosphatase inhibitors. Protein concentrations were determined using a bicinchoninic acid assay. Equal amounts of protein were separated by SDS-PAGE and transferred to PVDF membranes. Membranes were blocked in 5% skim milk in TBST and incubated overnight at 4°C with anti-RORγ antibody (Abcam; Cat. ab207082; 1:2000; approximately 55 kDa) and β-actin (Sigma-Aldrich; Cat. A5441; 1:10,000; 42 kDa) as a loading control. Horseradish peroxidase-conjugated secondary antibody (Bio-Rad; Cat. 1706515; 1:25,000) and chemiluminescent detection were subsequently used. Mouse thymus lysate was included as a positive control for RORγ expression.

For intracellular flow cytometry, YTN16-GFP and H59-GFP cells were seeded at 3 × $10^5$ cells per well and harvested at confluence using Accutase (BD Biosciences; cat 561527). Cells were stained with fixable eFluor 780 viability dye (ebioscience; Cat. 65-0865-14; 1:1000), fixed and permeabilized using the eBioscience FoxP3/Transcription Factor Staining Buffer Set and stained intracellularly with RORγ-PE (eBioscience; clone AFKJS-9; Cat. 12-6988-82; 1:50). Appropriate controls were used to establish RORγ positivity. Samples were acquired on a BD LSRFortessa and analysed using FlowJo v10.10.

For MC38-RFP cells, RORγ expression was assessed after 72 h exposure to vehicle or TF-S10 using the same immunofluorescence and intracellular flow-cytometric approaches. Relative RORγ-positive cell frequency was normalized to the corresponding untreated control where indicated.

#### S4.2 Direct tumour-cell response to TF-S10

Direct effects of TF-S10 on tumour-cell growth, viability, migration and apoptosis were assessed using clonogenic, metabolic viability, scratch-closure and Annexin V/propidium iodide assays.

For clonogenic assays, YTN16-GFP cells were seeded at 500 cells per well in collagen I-coated 24-well plates and allowed to adhere for 24 h. Cells were then treated with complete medium, vehicle or TF-S10 at 5, 10 or 25 μM and cultured for 7 days. Colonies were fixed and stained with 0.1% crystal violet in 40% ethanol, and total stained area was quantified using ImageJ.

For metabolic viability assays, YTN16-GFP cells were seeded at 500 cells per well in collagen I-coated 96-well plates. After 24 h, cells were treated with complete medium, vehicle or TF-S10 at 5, 10 or 25 μM. After 72 h, MTT reagent was added and metabolically active cells were quantified by absorbance at 565 nm. Viability was normalized to the corresponding control condition following background subtraction.

For scratch-closure assays, YTN16-GFP cells were seeded at $8 \times 10^4$ cells per well in collagen I-coated 24-well plates and grown to confluence. A linear scratch was generated using a sterile 200-μL pipette tip, detached cells were removed and cultures were treated with complete medium, vehicle or TF-S10 at 5 or 25 μM. Brightfield images were acquired immediately after scratch generation and after 72 h using an EVOS imaging system. Wound closure was quantified in ImageJ as the percentage reduction in wound width relative to baseline.

Apoptosis was assessed in MC38-RFP cells after 72 h exposure to complete medium, vehicle or TF-S10 at 5 or 25 μM. Cells were stained with Annexin V–Alexa Fluor 488 and propidium iodide using the Invitrogen Annexin V/Dead Cell Apoptosis Kit (Cat. V13241) and analysed by flow cytometry. Annexin V-positive/PI-negative cells were classified as apoptotic, whereas Annexin V-positive/PI-positive cells were classified as late apoptotic or necrotic.

### S5. Murine peripheral-blood and bone-marrow flow cytometry

#### S5.1 Peripheral-blood processing and immune phenotyping

Peripheral blood was collected into heparin-containing tubes at the time points specified for each experimental model. Red blood cells were lysed using 1× BD Pharm Lyse Lysing Buffer (Cat. 555899) for 10 min at room temperature in the dark. Lysis was repeated where required, with centrifugation at 450 × *g* for 5 min between steps. The resulting leukocyte fraction was stained with fixable eBioscience eFluor 780 viability dye (Cat. 65-0865-14; 1:1000) for 10 min on ice.

For surface immunophenotyping, cells were stained with Ly6G-FITC (clone 1A8; BioLegend, Cat. 127605), CD3-PE-Cy7 (clone 145-2C11; Invitrogen, Cat. 25-0031-82), CD4-PE-Cy5 (clone RM4-5; Invitrogen, Cat. 15-0042-82), CD8a-BV650 (clone 53-6.7; BD Horizon, Cat. 563234), CD11b-BV510 (clone M1/70; BioLegend, Cat. 101245) and PD-L1-APC (clone MIH5; Invitrogen, Cat. 17-5982-82). Surface antibodies were used at 1:200 unless otherwise indicated; PD-L1-APC was used at 1:50. Cells were incubated for 30 min on ice in the dark.

For intracellular staining, cells were fixed and permeabilized using the eBioscience FoxP3/Transcription Factor Staining Buffer Set (Cat. 00-5523-00) according to the manufacturer's instructions. Cells were stained with FOXP3-PE-eFluor 610 (clone FJK-16s; Invitrogen, Cat. 61-5773-82; 1:100), RORγ-PE (clone AFKJS-9; ebioscience, Cat. 12-6988-82; 1:50) and IL-17A-BV711 (clone TC11-18H10.1; BioLegend, Cat. 506941; 1:20) for 30 min at room temperature in the dark.

Single-stained compensation controls were prepared using antibody-capture beads or individually stained cells. Fluorescence-minus-one controls were used to establish gates where appropriate; for RORγ and IL-17A, the corresponding FMO controls incorporated matched isotype controls, whereas PD-L1 gating was based on FMO controls without an additional isotype control.

Samples were acquired on a BD LSRFortessa flow cytometer and analysed using FlowJo v10. Circulating neutrophils were identified as viable Ly6G$^+$CD11b$^+$ cells, with RORγ and other immune phenotypes assessed within the relevant gated populations.

#### S5.2 Bone-marrow processing and progenitor phenotyping

Femora and tibiae were harvested at the experimental endpoints specified for each model. Bone ends were removed and marrow was flushed using a 22-gauge needle with Hank's Balanced Salt Solution containing 0.5% fetal bovine serum and 20 mM HEPES. Marrow aggregates were mechanically dissociated to generate single-cell suspensions, centrifuged and subjected to erythrocyte lysis using 1× BD Pharm Lyse Lysing Buffer (Cat. 555899) for 10 min at room temperature.

Bone-marrow cells were stained with fixable eFluor 780 viability dye followed by a progenitor panel comprising lineage-eFluor 450 (Invitrogen, Cat. 88-7772-72; 1:5), CD34-Alexa Fluor 700 (clone 4H11; Invitrogen, Cat. 25-0349-41; 1:20), Sca-1-FITC (clone D7; Invitrogen, Cat. 11-5981-82; 1:200), CD117-APC (clone 2B8; Invitrogen, Cat. 17-1171-81; 1:143), CD16/32-PE-Cy7 (clone 93; Invitrogen, Cat. 25-0161-81; 1:77), CD135-PE (clone A2F10; Invitrogen, Cat. 12-1351-82; 1:20) and CD127-PE-Cy5 (clone A7R34; Invitrogen, Cat. 15-1271-82; 1:20).

Samples were acquired on a BD LSRFortessa and analysed using FlowJo v10. Progenitor populations were identified hierarchically within viable lineage-negative marrow cells. Conventional myeloid progenitors were defined within the Sca-1$^{-}$CD117$^{+}$ compartment, whereas emergency granulocyte–monocyte progenitors (eGMPs) were identified within the lineage-negative Sca-1$^{+}$CD117$^{+}$ compartment according to the gating strategy shown in **Figure 4**.

#### S5.3 Bone-marrow tissue processing for histology

For histological and spatial analyses, intact bones were fixed in 10% neutral-buffered formalin for 48 h and subsequently decalcified by the Histopathology Platform at the Research Institute of the McGill University Health Centre. Samples were paraffin embedded and stored at room temperature until sectioning and immunostaining.

### S6. Immunostaining of Murine Tissue Specimens

#### S6.1 Immunofluorescence staining of YTN16-GFP murine tissue specimens

Mouse lungs from the YTN16-GFP subcutaneous model were embedded in Tissue-Tek OCT compound (Sakura Finetek; cat. 4583), cryosectioned, fixed in 1% paraformaldehyde for 20 min at room temperature, permeabilized with 0.1% Triton X-100 for 1 min, and blocked for 1 h in TBST containing 0.01% Triton X-100, 2% BSA, and 1% FBS.

For metastatic burden analysis, sections were stained with DAPI (1:2000) for 15 min. For neutrophil analysis, sections were incubated with anti-mouse Ly6G-AF647 (BioLegend; cat. 127610; 1:50) and DAPI (1:1000) for 1 h at room temperature. Following PBS washes, sections were mounted with VECTASHIELD Vibrance Antifade Mounting Medium (Vector Laboratories; cat. H-1700-2) and imaged using a Zeiss LSM 780 confocal microscope. Images were analyzed in ImageJ. Metastatic burden was quantified as the percentage GFP-positive area, and neutrophils were quantified as $Ly6G^+DAPI^+$ cells. Statistical significance was determined using two-tailed Mann–Whitney tests ($P < 0.05$).

#### S6.2 Immunofluorescence staining of MC38-RFP murine tumour specimens

Multiplex immunofluorescence (mIF) was performed on formalin-fixed paraffin-embedded (FFPE) MC38-RFP tumour sections using the Opal™ tyramide signal amplification system (Akoya Biosciences) on an automated IntelliPATH FLX platform by the Molecular and Cellular Immunology Core at BC Cancer – Victoria Deeley Research Centre. The panel and staining conditions are summarized in **Supplementary Table S3**. Sections underwent sequential cycles of antigen retrieval, blocking, primary antibody incubation, HRP-mediated Opal fluorophore deposition, and antibody stripping, followed by DAPI counterstaining and mounting.

Whole-slide multispectral images were acquired on a Vectra Polaris system (Akoya Biosciences) at 20× magnification and analyzed using inForm software. Tissue and cell segmentation were performed using a supervised algorithm, and tissue was classified as tumour or non-tumour. Phenotyping algorithms (**Supplementary Table S5**) were applied to quantify distinct cell populations while preserving spatial coordinates. Quantitative cell counts and spatial analyses were exported using PhenoptrReports (Akoya Biosciences). Spatial analyses are described in **Supplemental Methods S7**.

#### S6.3 Immunohistochemistry staining of MC38-RFP murine lung and bone specimens

Multiplex immunohistochemistry (mIHC) was performed on formalin-fixed paraffin-embedded (FFPE) mouse lung and bone marrow sections by the Histopathology Core Facility, Research Institute of the McGill University Health Centre (RI-MUHC). Sections were stained on a Ventana automated platform using sequential multiplex staining with optimized antigen retrieval and antibody detection conditions. Whole-slide images were acquired using an Aperio digital slide scanner under standardized imaging conditions and analyzed using ImageJ. Details of the multiplex antibody panel and antibody dilutions are provided in **Supplementary Table S4**.

### S7. Cross-compartment spatial sampling and nested analysis

Spatial analyses were performed across tumour, bone marrow, and lung specimens to assess tissue-specific organization following RORγ inverse agonism. Because multiple local fields, tiles, or neighbourhoods were obtained from each biological specimen, these local spatial units were treated as nested subsamples rather than independent biological replicates. Primary treatment-group inference was therefore performed at the mouse or specimen level, with nested tile- or field-level models used as supporting analyses.

#### S7.1 Lung spatial analysis

Lung analyses included three vehicle-treated and three RORγ inverse agonist-treated mice. Cellular centroid coordinates derived from the image-analysis output were partitioned using a fixed Cartesian grid, with a primary tile size of 300 μm. Tiles containing fewer than 250 detected objects were excluded according to this fixed eligibility criterion. Local tiles were treated as spatial subsamples nested within mouse.

To assess robustness to grid placement, four prespecified grid origins were evaluated: (0,0), (0.5,0), (0,0.5), and (0.5,0.5) of the tile dimension. Additional sensitivity analyses used 250-, 400-, and 500-μm tile sizes. Tile assignment was determined geometrically from cellular centroid coordinates and was independent of treatment group or the visual appearance of the field.

For abundance endpoints, phenotype-positive fractions were calculated within each eligible tile. Within each grid origin, eligible-tile fractions were averaged without weighting to obtain a mouse-level value, and the four origin-specific estimates were subsequently averaged to generate the primary mouse-level summary.

For spatial co-association endpoints, Spearman correlations between phenotype fractions were calculated across eligible tiles within each mouse and grid origin. Correlation coefficients were Fisher-z transformed before averaging across grid origins and subsequently back-transformed for presentation.

Supporting nested analyses retained individual eligible tiles while accounting for their clustering within mouse. Abundance outcomes were transformed as arcsin[sqrt((positive count + 0.5)/(total object count + 1))]. For co-association analyses, phenotype fractions were independently converted to within-mouse midrank-normalized z scores and their product used as the local co-association outcome. Gaussian mixed-effects models included treatment as a fixed effect and mouse as a random intercept. Benjamini–Hochberg correction was applied across the prespecified family of 10 lung spatial outcomes within each scale/origin analysis.

#### S7.2 Bone-marrow spatial analysis

Bone-marrow analyses included two control and two RORγ inverse agonist specimens. Local cell neighbourhoods were defined using a 50-nearest-neighbour graph constructed from cellular spatial coordinates. The principal displayed neighbourhood endpoint was the fraction of cells located within neighbourhoods in which at least 50% of neighbouring cells were RORγ-positive.

A complementary tile-based analysis was performed as a sensitivity analysis and represented a distinct spatial estimand from the nearest-neighbour analysis. Each specimen was partitioned using a fixed 2,000 × 2,000 native-coordinate-unit grid anchored to the specimen-specific minimum centroid x and y coordinates. Tiles containing fewer than 100 objects were excluded using this fixed criterion. The resulting dataset contained 523 eligible tiles, comprising 248 control and 275 RORγ inverse agonist tiles.

For specimen-level tile summaries, the RORγ-positive fraction was calculated within each specimen from the summed number of RORγ-positive cells divided by the summed number of all cells across eligible tiles. Individual tiles were not treated as independent biological replicates. Supporting nested analyses used arcsin[sqrt((positive count + 0.5)/(total object count + 1))]-transformed tile fractions with specimen-level clustering accounted for using a specimen random-intercept model and a specimen-cluster-robust sensitivity analysis.

The ≥50% RORγ-positive neighbourhood endpoint and the 2,000-unit tile RORγ-positive fraction therefore quantify different spatial properties and were interpreted separately.

#### S7.3 Tumour spatial analysis

Tumour spatial analyses included three control and three RORγ inverse agonist-treated mice, with annotation fields treated as spatial subsamples nested within mouse. Quantitative cell identities, phenotype calls and global centroid coordinates were obtained from the retained consolidated vendor object export. These upstream segmentation and phenotype classifications were treated as locked source data and were not regenerated or re-thresholded during the present analysis.

Endpoint-specific field eligibility criteria were applied independently of treatment effect. RORγ/PD-L1 intensity analyses required ≥1,000 tumour objects, ≥5 objects of the relevant lineage and a finite positive field median. Analyses of the RORγ-positive and PD-L1-positive myeloid fractions required ≥1,000 tumour objects and ≥1 myeloid object. Directional spatial analyses required ≥1,000 tumour objects and ≥20 coordinate-valid myeloid/CD8 objects.

Ly6G-rich hotspot fields were defined as sampled tumour fields in which Ly6G$^{+}$ cells comprised >1% of all evaluable cells with a binary Ly6G call. Additional descriptive thresholds of >5% and >10% Ly6G$^{+}$ cells were examined to assess progressively higher-density Ly6G enrichment. For the >1% threshold, the proportion of Ly6G-rich fields was calculated within each mouse and treatment groups were compared using an exact mouse-level test. Across all sampled fields, 9/30 control fields and 2/30 RORγ inverse agonist-treated fields met the >1% threshold; these pooled field counts were displayed descriptively, while statistical inference retained mouse as the biological replicate.

Ly6G–CD8 spatial interactions were quantified as the number of Ly6G–CD8 cell pairs separated by ≤25 μm within eligible tumour fields. Pair counts were summed within each mouse to generate the mouse-level values displayed in the figure. The corresponding totals across mice were 30,141 pairs in control tumours and 720 pairs in RORγ inverse agonist-treated tumours, representing a 97.6% descriptive reduction. Treatment groups were compared using an exact mouse-level test. Because pair counts were strongly right-skewed and included zero values, mouse-level values were displayed after $\log_{10}$(count + 1) transformation for visualization only; statistical inference was based on the prespecified mouse-level comparison. Field-level mixed-effects log-count models, including maximum-likelihood and restricted maximum-likelihood fits, together with a mouse-clustered robust analysis, were treated as exploratory supporting analyses of the nested field-level observations.

For other tumour spatial endpoints, eligible field values were averaged without weighting within each mouse to obtain one biological value per mouse for primary treatment-group inference. Between-group comparisons therefore retained mouse, rather than annotation field, as the biological replicate. Field-level mixed-effects and mouse-clustered analyses were used only as supporting analyses of the nested spatial observations.

#### S7.4 Tumour representative-image reconstruction and display

Representative tumour microscopy images were reconstructed from the original PerkinElmer QPI/QPTIFF files and the retained consolidated vendor object export. Global vendor cell-centroid coordinates were transformed independently for each slide into native QPTIFF pixel coordinates using the slide-specific TIFF position metadata and the exact native calibration of 0.4966789857709499 μm/pixel. Original phenotype and coordinate fields were preserved, and no cell or nuclear polygons were inferred.

Exact vendor FlaggedForAnalysis rectangles and deterministic RORγ-positive and RORγ-negative registration-QC centroid points were validated in QuPath 0.6.0. QuPath was used for native image access, registration validation, storage of QC objects, image export and overlay rendering; it was not used to re-segment cells or modify retained phenotype classifications.

Each tumour slide contained ten FlaggedForAnalysis rectangles. One representative rectangle per mouse was selected using a deterministic within-slide typicality rule: the rectangle whose RORγ-positive fraction was closest to the median of the ten rectangles was selected, with ties resolved by larger classified-cell count and subsequently by annotation identifier. Treatment group and visual appearance were not used in this ranking.

Native-resolution representative images were exported using fixed linear 0–255 channel displays without per-image normalization. For figure readability, display-only companions were generated using a single pooled linear display window applied identically across all six mice, with output values calculated as min[255, round(input × 255/64)]. No gamma transformation, histogram equalization, background subtraction, denoising, sharpening, deconvolution, treatment-specific adjustment, generative image processing or manual pixel editing was used.

Centroid overlays represent the positions of retained vendor phenotype calls and do not represent cellular boundaries or segmentation contours. All tumour quantification, spatial distances and statistical analyses were derived from the locked object and coordinate data and not from the display PNGs.

#### S7.5 Inferential hierarchy and avoidance of pseudoreplication

Across all three compartments, tiles, annotation fields, and cellular neighbourhoods represented local spatial subsamples of the originating mouse or specimen and were not counted as independent biological replicates. Primary group-level inference was based on mouse- or specimen-level summaries. When local spatial observations were analysed directly, mixed-effects or cluster-aware models were used to account for within-replicate dependence. These nested analyses were interpreted as supporting evidence of within-tissue spatial organization rather than as additional biological replication.

### SUPPLEMENTAL TABLES

**Supplementary Table S1. Demographic and Clinicopathologic Characteristics of The Peripheral Blood Flow Cytometry Patient Cohort and Tumour Tissue Microarray Cohort.**

**Abbreviations**: PB: peripheral blood; cNLR: circulating neutrophil-to-lymphocyte ratio; TMA: tumour microarray

| Characteristic | Total PB flow cohort (n=26) | Low cNLR PB Cohort (n=18) | High cNLR PB Cohort(n=8) | NYC TMA (n=29) |
|---|---|---|---|---|
| **Age, median (IQR), y** | 68 (61.3–75.8) | 70 (60.3–76.8) | 67 (62.8–70.5) | 66 (60–70) |
| **Male sex, n/N (%)** | 21/26 (80.8%) | 13/18 (72.2%) | 8/8 (100%) | 26/29 (89.7%) |
| **cNLR, median (IQR)** | 3.03 (2.16–4.23) | 2.38 (1.91–3.05) | 4.91 (4.61–5.20) | 2.44 (2.16–5.52) |
| **Stage I, n/N (%)** | 5/26 (19.2%) | 4/18 (22.2%) | 1/8 (12.5%) | 2/29 (6.9%) |
| **Stage II, n/N (%)** | 6/26 (23.1%) | 5/18 (27.8%) | 1/8 (12.5%) | 7/29 (24.1%) |
| **Stage III, n/N (%)** | 9/26 (34.6%) | 6/18 (33.3%) | 3/8 (37.5%) | 18/29 (62.1%) |
| **Stage IV, n/N (%)** | 6/26 (23.1%) | 3/18 (16.7%) | 3/8 (37.5%) | 2/29 (6.9%) |
| **Neoadjuvant therapy, Y / N / N/A** | 15 / 7 / 4 | 10 / 6 / 2 | 5 / 1 / 2 | 20 / 9 / 0 |

**Supplementary Table S2. Summary of Antibodies Used for 6-plex Immunostaining of Human Tumour Microarrays.** All the OPAL TSA fluors listed here are from Akoya Biosciences. **Abbreviations:** Rb = rabbit; Ms = mouse; CK7 = Cytokeratin 7; Arg1b = Arginase 1b

| Antigen (clone) | Vendor, Cat # | Retrieval | Dilution Factor | Incubation [min] | Secondary or 2o HRP Polymer | Vendor/ Cat # | Incubation [min] | Opal Fluorophore | Cat# Fluor | Fluorophore Dilution | Incubation [min] |
|---|---|---|---|---|---|---|---|---|---|---|---|
| IL-17A (poly) | R&D, AF-317-NA | ER2-20 | 100 | 30 | Goat HRP polymer 2 step | Biocare/ GHP516 | 10+10 | 690 | FP1497001KT | 150 | 10 |
| RORγ (6F3.1) | Millipore, MABF81 | ER2-20 | 300 | 30 | Akoya HRP Polymer (Rb + Ms) | Akoya/ ARH1001EA | 10 | 480 | FP1500001KT | 150 | 10 |
| CXCL12 (poly) | Abcam, ab9797 | ER2-20 | 400 | 30 | Akoya HRP Polymer (Rb + Ms) | Akoya/ ARH1001EA | 10 | 570 | FP1488001KT | 150 | 10 |
| CXCR4 (UMB2) | Abcam, ab124824 | ER2-20 | 500 | 30 | Akoya HRP Polymer (Rb + Ms) | Akoya/ ARH1001EA | 10 | 520 | FP1487001KT | 150 | 10 |
| Arg1b (Poly) | Genetex, GTX109242 | ER1-20 | 150 | 30 | Akoya HRP Polymer (Rb + Ms) | Akoya/ ARH1001EA | 10 | 620 | FP1495001KT | 150 | 10 |
| CK7 (EPR17078) | Abcam, ab181598 | ER2-20 | 4000 | 30 | Akoya HRP Polymer (Rb + Ms) | Akoya/ ARH1001EA | 10 | 780 | FP1501001KT | 25 | 60 |

**Supplementary Table S3: Summary of Antibodies Used for 7-plex Immunostaining of Murine Tumour Samples.** The antibodies are listed in the order in which they were sequentially stained in. All the fluors listed here are from Akoya Biosciences. Abbreviations: Rb = rabbit; TSA-DIG = Tyramide Signal Amplification – Digoxigenin

| Round | $1^0$ Ab | Species | Cat# | Clone | AR Before | Block | Dilution | Polymer | OPAL Fluor | Fluor Cat# |
|---|---|---|---|---|---|---|---|---|---|---|
| **1** | RORg | Rb | ab207082 | EPR20006 | Nuclear | Terminator 12, no wash | 1/500 | M2 Rb HRP | 620 | FP1495001KT |
| **2** | CD163 | Rb | ab182422 | EPR19518 | AR9 | Sniper 10 | 1/2000 | M2 Rb HRP | 690 | FP1497001KT |
| **3** | PanCK | Rb | ab308262 | EPR28285-45 | AR9 | Sniper 10 | 1/5500 | M2 Rb HRP | 570 | FP1488001KT |
| **4** | Ly6G | Rb | 87048S | E6Z1T | AR6 | Sniper 10 | 1/100 | M2 Rb HRP | 480 | FP1500001KT |
| **5** | PD-L1 | Rb | 80517 | F3ZT1T | AR9 | Sniper 10 | 1/100 | M2 Rb HRP | 520 | FP1487001KT |
| **6** | CD8 alpha | Rb | 98941S | D4W2Z | AR6 | Sniper 10 | 1/100 | M2 Rb HRP | TSA-DIG | N/A |
| **7** | 780 | - | - | - | AR9 | - | 1/25 | - | - | N/A |

**Supplementary Table S4: Summary of Immunostaining Protocol for Murine Tissues. Abbreviations**: IF: immunofluorescence; IHC: immunohistochemistry; anti-ms: anti-mouse

| Samples | Fixation/Frozen | Antibodies (dilution) | Cat# | Imaging/Scanning | Analysis (Technique) |
|---|---|---|---|---|---|
| **Lungs (YTN16-GFP cohort)** | Optimal Cutting Temperature (Sakura Finetek) | Ly6G-FITC (1:50) | Biolegend; cat 127610 | Zeiss LSM 780 | ImageJ (IF) |
| **Bones (MC38-RFP cohort)** | 10% Formalin | Anti-ms RORγ (1:100) | LS-Bio: LS-A5191-50 | Ventana/Aperio | HALO (IHC) |
| **Lungs (MC38-RFP cohort)** | 10% Formalin | Anti-ms CD8 (1:50)<br>Anti-ms Ly6G (1:100)<br>Anti-ms CD163 (1:400)<br>Anti-ms F4/80 (1:50) | Cell Signaling, 98941<br>BD Biosciences; 551459<br>Abcam; ab182422<br>Invitrogen; 14-4801-82 | Ventana/Aperio | HALO (IHC) |
| **Tumours (MC38-RFP cohort)** | 10% Formalin | Refer to suppl. Table S3 | Refer to suppl. Table S3 | Phenomalger/Vectra Polaris (Akoya Biosciences) | InForm/QuPath (OPAL IF) |

**Supplementary Table S5: Phenotype Models Used for Mouse Tumour Sample Analysis.** The 12 phenotypes were used for the analysis of 6-plex immunostaining of mouse tumour samples. Red denotes negativity for the marker, while green denotes positivity for the respective marker. Abbreviations: PanCK = Pan Cytokeratin; PD-L1 = Programmed Death Ligand 1

| | **CD8** | **CD163** | **Ly6G** | **PanCK** | **RORγ** | **PD-L1** |
|---|---|---|---|---|---|---|
| **Cancer cells** | Red | Red | Red | Green | Red | Red |
| **CD8+ T cells** | Green | Red | Red | Red | Red | Red |
| **M2 Macrophages** | Red | Green | Red | Red | Red | Red |
| **RORγ+ M2 Macrophages** | Red | Green | Red | Red | Green | Red |
| **PD-L1+ M2 Macrophages** | Red | Green | Red | Red | Red | Green |
| **Neutrophils** | Red | Red | Green | Red | Red | Red |
| **PDL1+ neutrophils** | Red | Red | Green | Red | Red | Green |
| **RORγ+ neutrophils** | Red | Red | Green | Red | Red | Green |
| **Tumour PD-L1** | Red | Red | Red | Green | Red | Green |
| **Tumour RORγ** | Red | Red | Red | Green | Green | Red |
| **RORγ+ CD8+ T cells** | Green | Red | Red | Red | Green | Red |
| **PD-L1+ CD8+ T cells** | Green | Red | Red | Red | Red | Green |

### SUPPLEMENTARY FIGURES

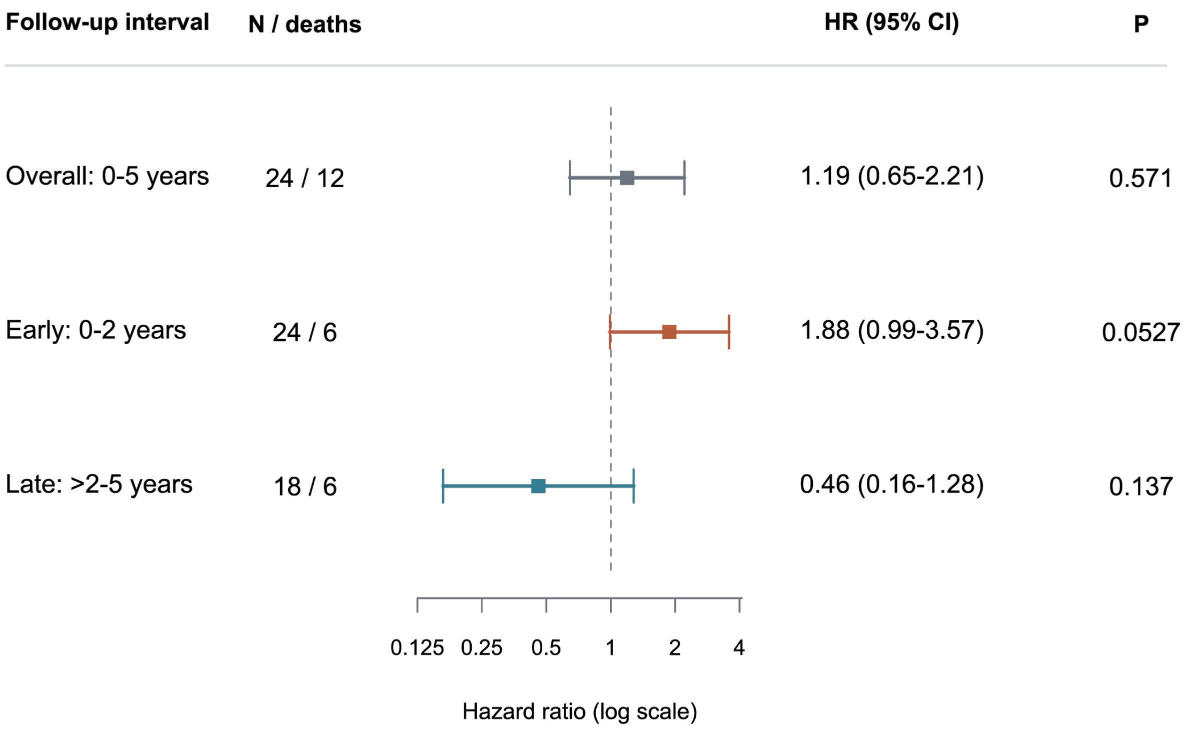


**Supplementary Figure S1. Tumour-Core IL-17+ clustering and post-operative survival.** Exploratory continuous Cox models │ hazard ratio per baseline sample SD. Time-period interaction: nominal likelihood-ratio P = 0.0225. The two-year boundary was selected post hoc. Late estimates are conditional on survival and follow-up beyond two years.

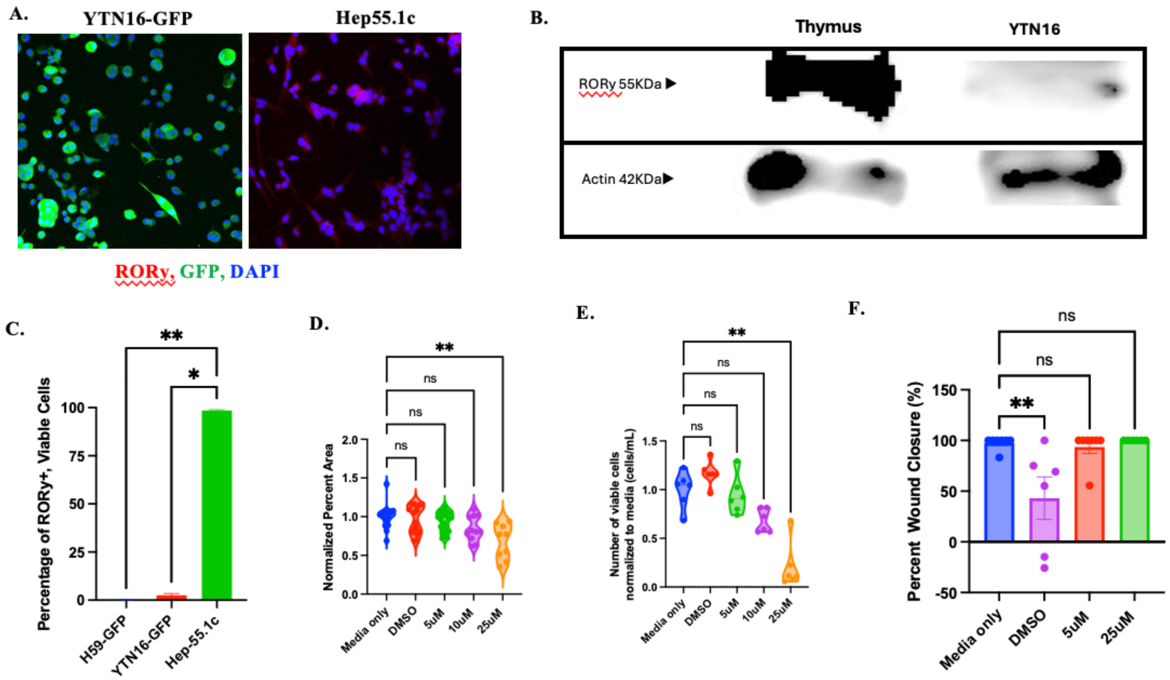


**Supplementary Figure S2: TF-S10 does not affect cell growth and viability in low-RORγ expressing YTN16 cancer cell line.** (A) Immunofluorescence staining of RORγ expression in YTN16-GFP cancer cell line and Hep55.1c cancer cell line. (B) Western blot showing RORγ expression in YTN16 cancer cell line. (C) Flow cytometric analysis of RORγ expression in multiple murine cancer cell lines. (D) YTN16 colony formation assay under different conditions. (E) YTN16 cell viability assay under different conditions. (F) YTN16 scratch assay under different conditions.

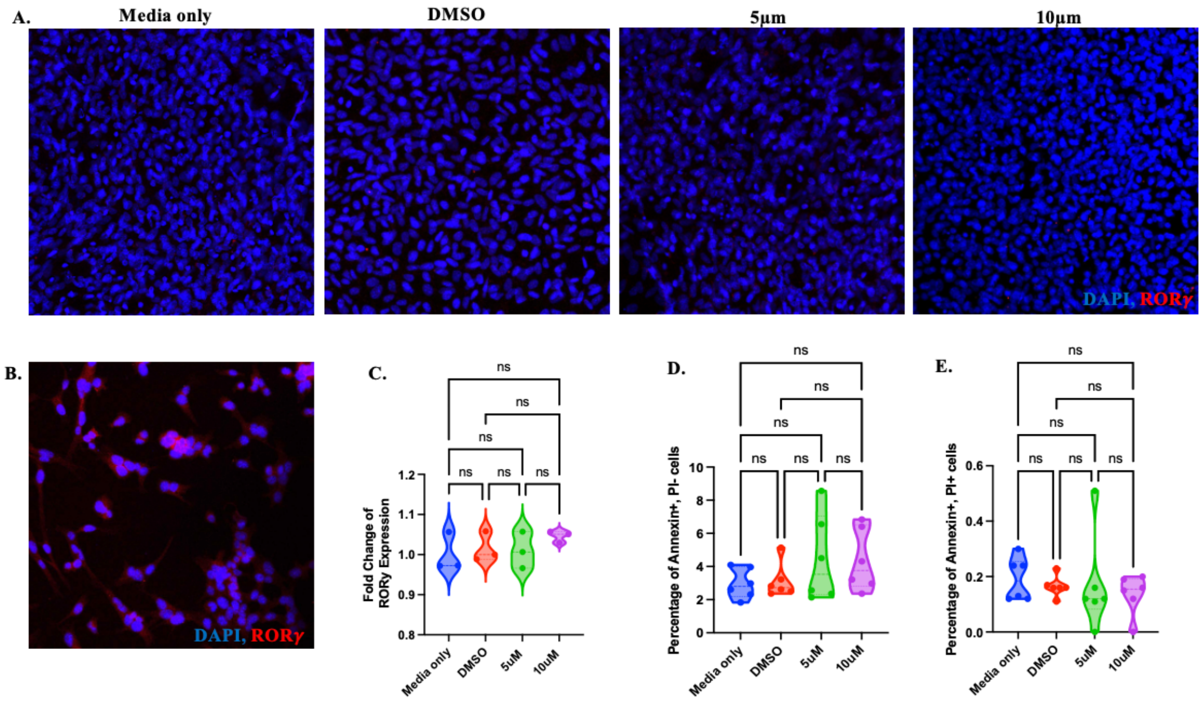


**Supplementary Figure S3: TF-S10 does not affect RORγ expression and cell viability in MC38 cancer cell line.** (A) Immunofluorescence staining of RORγ expression in MC38 cancer cell line under different conditions. (B) Immunofluorescence staining of RORγ expression in a positive control Hep55.1c cancer cell line. (C) Flow cytometric analysis of fold change RORγ expression in MC38 cell line under different conditions. (D-E) Prevalence of (D) apoptotic and (E) necrotic MC38 cells under different treatment conditions.

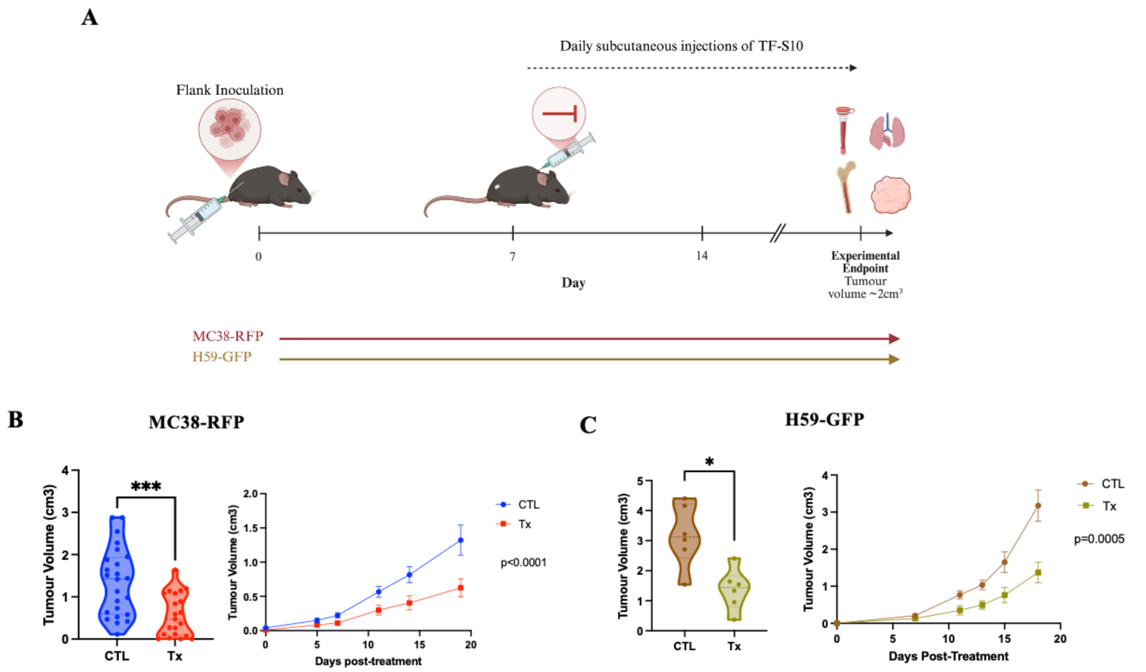


**Supplementary Figure S4: RORγ inverse agonism suppresses established tumour progression across different murine models.** (A) Experimental design. Syngeneic MC38-RFP colorectal and H59-GFP lung tumours were established subcutaneously in C57BL/6 mice. RORγ inverse agonist TF-S10 (Tx; MC38-RFP, n=21; H59-GFP, n=6) and vehicle (CTL; MC38-RFP, n=24; H59-GFP, n=6) treatment commenced on day 7 after tumour implantation and continued until experimental endpoint. (B–C) Longitudinal tumour growth curves and endpoint tumour volume demonstrating significant suppression of established tumour progression in (B) MC38-RFP colorectal and (C) H59-GFP lung tumour models under different treatment conditions.
